# Considering dynamic nature of the brain: the clinical importance of connectivity variability in machine learning classification and prediction

**DOI:** 10.1101/2023.01.26.525765

**Authors:** Inuk Song, Tae-Ho Lee

## Abstract

The brain connectivity of resting-state fMRI (rs-fMRI) represents an intrinsic state of brain architecture, and it has been used as a useful neural marker for detecting psychiatric conditions as well as for predicting psychosocial characteristics. However, most studies using brain connectivity have focused more on the strength of functional connectivity over time (static-FC) but less attention to temporal characteristics of connectivity changes (FC-variability). The primary goal of the current study was to investigate the effectiveness of using the FC-variability in classifying an individual’s pathological characteristics from others and predicting psychosocial characteristics. In addition, the current study aimed to prove that benefits of the FC-variability are reliable across various analysis procedures. To this end, three open public large resting-state fMRI datasets including individuals with Autism Spectrum Disorder (ABIDE; *N* = 1249), Schizophrenia disorder (COBRE; *N* = 145), and typical development (NKI; *N* = 672) were utilized for the machine learning (ML) classification and prediction based on their static-FC and the FC-variability metrics. To confirm the robustness of FC-variability utility, we benchmarked the ML classification and prediction with various brain parcellations and sliding window parameters. As a result, we found that the ML performances were significantly improved when the ML included FC-variability features in classifying pathological populations from controls (e.g., individuals with autism spectrum disorder vs. typical development) and predicting psychiatric severity (e.g., score of autism diagnostic observation schedule), regardless of parcellation selection and sliding window size. Additionally, the ML performance deterioration was significantly prevented with FC-variability features when excessive features were inputted into the ML models, yielding more reliable results. In conclusion, the current finding proved the usefulness of the FC-variability and its reliability.

## Introduction

The resting-state based functional connectivity is considered as a representation of the individual’s stable neural foundation that determines how individual responds to task-specific demands or environments (Cole et al., 2014; He, 2013; Smith et al., 2013). Indeed, this intrinsic functional connectivity pattern carries considerable promise as a tool in providing potential neural markers for various range of human behaviors including mental disorders as everyone has their own unique intrinsic patterns with a fairly high stability across time (Cole et al., 2014; Finn et al., 2015; Noble et al., 2017; Taxali et al., 2021; Termenon et al., 2016). At the early stage of human functional connectivity (FC) studies, researchers commonly adopted static measures of connectivity (static-FC), in which pairwise correlation (e.g., Pearson correlation) between time-series of each brain regions is calculated. However, as the static method uses entire BOLD signals in connectivity calculation, it does not consider temporal dynamics of neural communications that spontaneously changes over time (Chang & Glover, 2010; Chang et al., 2013). However, recent studies have demonstrated that the spontaneous signal changes significantly linked to the physiological underpinnings of information processing in the brain (Allen et al., 2018; Matsui et al., 2019; Tagliazucchi et al., 2012; Thompson et al., 2013) such that that temporal dynamics of FC in hemodynamics is significantly correlated with the calcium-imaging based FC measure which is a more direct measure of neuronal activations (Matsui et al., 2019).

Recently, time-varying or dynamic-FC has gained attention to consider the temporal characteristics in estimation of functional connectivity as a promising substitute (Calhoun et al., 2014; Cohen, 2018; Hutchison et al., 2013; Preti et al., 2017). Typically, the dynamic-FC is estimated by using a sliding-window method which calculate FC within a smaller time window, the window slides to the next time, and this procedure is repeated, yielding many subsets of connectivity patterns. Thus, the dynamic approach can preserve the temporal dynamic information of connectivity across time. The derived dynamic-FC can be used in two approaches mostly to identify temporal features of FC (Fong et al., 2019). The ‘state-based’ approach distinguishes several dynamic states by identifying repeated spatial dynamic-FC profiles using clustering algorithms such as k-means clustering (Weng et al., 2020) and evolutionary clustering (Deshpande & Jia, 2020). In contrast, the ‘edge-based’ approach which is more focused on the temporal features of each dynamic-FC (Preti et al., 2017), indicating connectivity stability over time quantified by using standard deviation (Chen et al., 2017; Falahpour et al., 2016; Kucyi & Davis, 2014; Kucyi et al., 2013; Li et al., 2018), concordance coefficient (L. Li et al., 2020), and mean square successive difference (MSSD; Chen et al., 2021).

Recent functional connectivity studies using the ‘edge-based’ approach have highlighted that the temporal variability characteristic, namely functional connectivity variability (FC-variability; Elton & Gao, 2015; Kucyi & Davis, 2014), can be more sensitive in classifying disorders compared to the static approach. For example, studies of consciousness with FC-variability showed that spontaneous FC patterns are related to the stream of cognitive processes rather than random fluctuations which was shaped by anatomical connectivity in anesthetized animals (Barttfeld et al., 2015; Hudetz et al., 2015; Hutchison et al., 2014) and humans (Kucyi & Davis, 2014). In psychiatric disorders, it was revealed that different FC-variability patterns for autism spectrum disorder (Chen et al., 2017; Chen et al., 2021; Falahpour et al., 2016; Guo et al., 2020; Harlalka et al., 2019; Kim et al., 2021; Y. Li et al., 2020; Mash et al., 2019), epilepsy (Li et al., 2018), schizophrenia (Supekar et al., 2019), ADHD (Wang et al., 2018), and depressive disorders (Chen et al., 2022). The FC-variability has further yielded cognitive applications in attention and arousal of caffeine intake. The attentional ability was associated with more stable FC-variability between brain networks (Fong et al., 2019) and the higher arousal after caffeine dose was represented as increasing in FC-variability (Rack-Gomer & Liu, 2012).

However, the benefits of FC-variability have not been studied enough. For instance, to examine the benefits, one thing needed is to compare static-FC and FC-variability. Only a few studies have compared static-FC and FC-variability (Fong et al., 2019; Wang et al., 2018). Since the studies only focused on specific functional domains such as attention, it is still unclear whether FC-variability can contribute to explaining other cognitive functions. Furthermore, previous studies have used different procedures such as neuroimage preprocessing, defining brain regions (i.e., brain parcellation selection), and dynamic-FC calculation. As shown in the previous study, parcellation selection can affect static-FC and dynamic-FC calculation (Bryce et al., 2021), and different time window sizes of the sliding-window approach also can affect the calculated dynamic-FC (Zalesky & Breakspear, 2015). It is necessary to examine beneficial effect of the FC-variability in multiple functional domains using a consistent procedure.

In this study, we systematically investigated the benefits of FC-variability across various domains such as group classification and individuals’ characteristics prediction using three large open-public rs-fMRI datasets: ABIDE, COBRE, and NKI (see Methods section). The robust benefits of FC-variability inclusion in the group classification and prediction model were also demonstrated by varying analytic procedures including different brain parcellation schemes at both regional and network levels including AAL2 (Rolls et al., 2015), Schaefer200 (Schaefer et al., 2018), and LAIRD (Laird et al., 2011) atlases and sliding-window sizes from 60 sec. to 120 sec. In addition, that the classification and prediction results would reveal additional important brain regions which have been overlooked when the static-FC was used solely.

## Methods

### Resting-state fMRI data

Resting-state functional MRI data were obtained from publicly available datasets including the Autism Brain Imaging Data Exchange I & II (ABIDE; Di Martino et al., 2017; Di Martino et al., 2014), the Center for Biomedical Research Excellence (COBRE^1^) and the enhanced Nathan Kline Institute (NKI)-Rockland (Nooner et al., 2012). The datasets were initially downloaded through the Mind Research Network’s collaborative informatics and neuroimaging suite (COINS; Landis et al., 2016). In the main analyses, this study only included individuals with T1 structural images, more than 100 volumes in echo-planar imaging (EPI) images, full-coverage of cerebral cortex, and without severe head motions (FD < 0.25 mm). In ABIDE II, the longitudinal collections (UCLA_long, UPSM_long) were excluded because the same subjects participated in the ABIDE I. As a result, the ABIDE consisted of 537 autism spectrum disorder (ASD) individuals (age: 15.9 ± 8.2, 75 females) and 712 typical development (TD; 15.4 ± 7.8, 185 females). The COBRE consisted of 71 schizophrenia (38.1 ± 13.9, 14 females) and 74 control (35.8 ± 11.6, 23 females). The NKI, a developmental dataset across the lifespan had 672 individuals ranging from 6 to 83 years of age (40.5 ± 20.6, 429 females). There were no significant differences in age within each dataset (ABIDE: *p* = 0.22, COBRE: *p* = 0.28).

Given that previous findings that the connectivity estimation is not markedly affected by sampling rates (Huotari et al., 2019; Nomi et al., 2017), the BOLD timeseries of the ASD dataset (ABIDE) was resampled to 0.33 Hz (i.e., TR = 3000 ms; Huotari et al., 2019) to set the same sliding-window size of dynamic-FC estimation as the ABIDE was collected with various sampling rates^2^. The schizophrenia dataset (COBRE) and the developmental dataset (NKI) had identical sampling rates between group samples (e.g., ASD vs. TD), 0.5 Hz and 0.7 Hz respectively, the time-series for those datasets was not resampled. Additional demographic information is available at https://osf.io/xd3fe/.

### Preprocessing

Structural and functional images were preprocessed using the FMRIB software library (FSL; Jenkinson et al., 2012) with ICA-AROMA (Pruim et al., 2015). The structural image was skull-stripped and segmented to tissue mask (CSF/WM/GM) after bias-field correction. The functional data were preprocessed including first ten volumes cut, motion correction, 5-mm smoothing, slice-timing correction, intensity normalization, ICA denoising (corrected mean FD; ABIDE: 0.025 mm, COBRE: 0.182 mm, NKI: 0.025 mm), band-pass filtering (0.001 – 0.08 Hz) and regressing out CSF/WM signal. The functional images were then normalized to the standard MNI 2-mm brain template through the non-linear transformation using the ANTs (Avants et al., 2014).

### Functional connectivity for static and dynamic levels

In the main functional connectivity analysis, BOLD time-series was first extracted from all regions using a structure-based atlas (AAL2; Rolls et al., 2015), a rs-fMRI-based atlas (Schaefer200 with Yeo 7networks; Schaefer et al., 2018), and a networklevel atlas (Laird et al., 2011). Several functional data that does not fully cover cerebellum regions in the data acquisition, and thus the cerebellum ROIs were excluded for further analyses, resulting in 94 AAL2 ROIs. Schaefer200 atlas covered cerebral areas. Similarly, ‘noise’ networks from LAIRD were excluded resulting in 18 networks.

For the static-FC estimation, robust correlation method was adopted using *‘robustfit’* function in MATLAB 2018a with the iterative reweighted least squares to minimize the potential impact of outliers in the connectivity estimation (Choi et al., 2012; Kinnison et al., 2012; Shen et al., 2017). By selecting two different brain regions iteratively, a set of pairwise connectivity was estimated. Finally, the estimated connectivity was then Fisher z-transformed for subsequent machine learning classification and prediction.

For the dynamic-FC and subsequent FC-variability estimation (i.e., edge-based approach), we used a sliding-window approach with robust correlation. FCs were calculated for each 60s-sliding window. To quantify the degree of variability, the mean square successive difference approach (MSSD; Von Neumann et al., 1941) was adopted to minimize the influence of gradual shifts of fMRI signal (Uddin, 2020). The MSSD was calculated by subtracting the dynamic-FC value of a given sliding window at t from t+1, squaring the result and finally averaging the squared values (see equation 1). The MSSDs were z-score normalized.

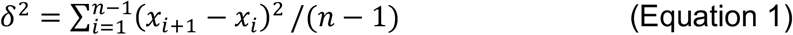

Although there is no gold standard in choice of sliding window size and previous studies used different size (e.g., 30s - 100s; Please see Fig S1 in Leonardi & Van De Ville, 2015; Preti et al., 2017; Shirer et al., 2012; Zalesky & Breakspear, 2015), 60s size was selected here as it is between the most commonly used window lengths. In the supplementary document, we also provided additional results repeated with different sizes of sliding window (90s and 120s) to confirm the robustness of the current findings. The results using the longer window sizes were consistent with the results of main finding (see Supplementary Table 1-4).

### Feature selection

The FC estimations at both static and dynamic levels yielded n×n matrices of overall connectivity strength over time (i.e., static-FC) and FC-variability (i.e., MSSD), where n is the number of ROIs (e.g., 94 for the AAL2). Since the matrices were diagonally symmetric, only lower triangle part of the matrices was used for further analyses.

The static-FC and/or FC-variability was then put into lasso regression for selecting informative features (Kassraian-Fard et al., 2016). In the absence of the selected features, when predicting PANSS scores, elastic-net (α = 0.5) was used to secure more features. There were three models and subsequent machine learning (ML) results were compared: 1) static-FC, 2) FC-variability, and 3) appended the static-FC and the FC-variability (i.e., combined).

A lambda parameter, the strength of regularization, was determined using 200 grid-search with 5-fold cross-validation with 10 iterations using the ‘lassoglm’ function in MATLAB. Still, there is a possibility of missing informative features from the lasso regression, we first used the ‘smallest lambda’ of 200 grid-search (i.e., least stringent criteria). Static-FC and/or FC-variability features whose non-zero lasso regression coefficients were inputted into the subsequent support vector machine (SVM) or support vector regression (SVR) in the order of magnitude of the coefficients. This procedure allows to control the number of features as well as estimate the benefits of FC-variability overall.

In addition, the ‘optimal’ lambda was also determined as the mean across 10 iteratively calculated lambda values which minimized the deviance of 5-fold crossvalidation (Teipel et al., 2017) to demonstrate the benefits when specific lambda parameters were selected. Similarly, after the optimal lambda was set, the features whose non-zero coefficients were used to train subsequent machines.

### Machine learning classification and prediction

The SVM was adopted to classify the ASD and the TD in the ABIDE and the schizophrenia and the control in the COBRE. The SVR was used to predict individuals’ characteristics. Since not all participants had available scores, subsets of datasets were used for prediction. We used representative variables to predict from the datasets: For the ASD, autism diagnostic observation schedule (ADOS; Lord et al., 2000, *N* = 377); For the schizophrenia, total positive and negative scores from positive and negative syndrome scale (PANSS; Kay et al., 1987, *N* = 69); For the NKI dataset, participants’ age was used (*N* = 672; see Supplementary Figure 2). The ML analyses were conducted by using MVPA-Light toolbox (Treder, 2020) and LIBSVM (Chang & Lin, 2011).

The SVM or SVR results were evaluated by 5-fold cross-validation with 100 iterations. To evaluate the overall performance of SVM, the area under curve (AUC) metric was used since the metric is independent of changes to the classifier threshold. For the SVR predictions, the mean squared error (MSE) was used to evaluate prediction performance. Since the main results using the smallest lambda produced line graphs as a function of number of features, secondary AUC from the line graphs were calculated for statistical testing. The statistical tests to compare the secondary AUCs of static-FC, MSSD, and the combined were performed using permutation test with 5000 iterations with Bonferroni correction. The classification results using the optimal lambda in the feature selection were also tested. Example codes are available at https://osf.io/xd3fe/.

## Results

### Group classifications

As shown in Figure 2, the additional MSSD features (i.e., ‘combined’) increased SVM classification performances overall. The line graphs show overall classification performances with the smallest lambda in the feature selection stage, and the bar graphs show corresponding secondary AUCs. Except for one classification between the schizophrenia and the control using the AAL2 atlas, all classification performances demonstrated benefits of including FC-variability (all *p* < .001; Table 1). Even in the exceptional case, the secondary AUC was almost identical (static-FC: 0.967, combined: 0.965). Since the static-FC already had a high classification performance, perhaps there was no room for further performance increase. Furthermore, the results implied that including FC-variability caused more reliable classification performance. If too many features which might include less-informative features were inputted into the SVM, the classification performances tended to worse due to the features’ complexity when the static-FC or MSSD features were used solely. In contrast, the performances of the ‘combined’ were increased reliably. The increased classification performances by including FC-variability were consistent when the optimal lambda was used in the feature selection (Supplementary Figure 1). We also identified some features of FC-variability that contributed to classifying the groups by scrutinizing the weights of SVM (Supplementary Figure 3, 4).

**Figure 1.**
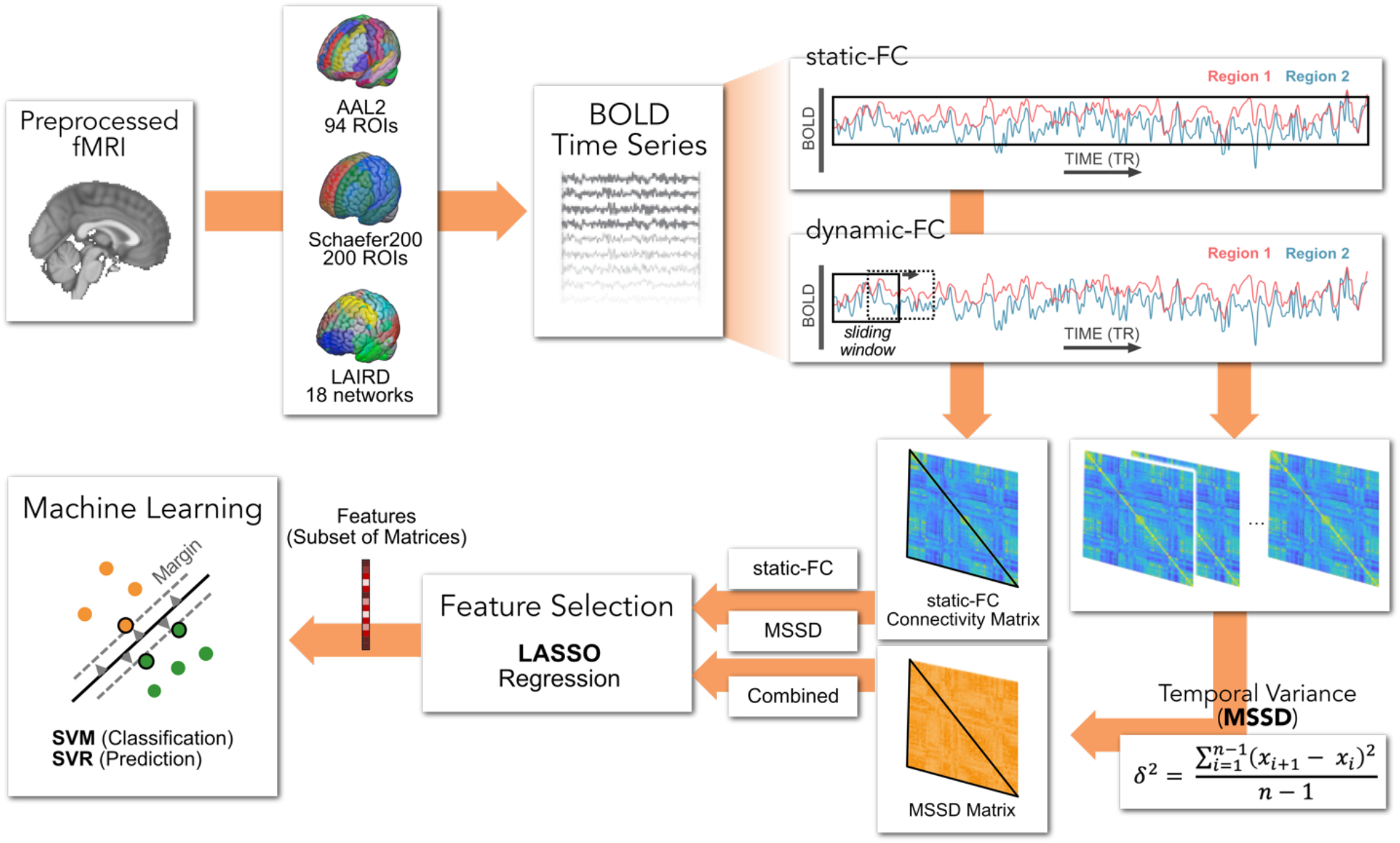
Overview of the analytical process in the current study. The BOLD signals were extracted from resting-state fMRI using AAL2, Schaefer200 atlases, and LAIRD network atlas. The signals were then used to calculate static-FC and moment-to-moment variability of dynamic-FC (MSSD). After the lasso regression to select a subset of components (feature selection), the selected features were utilized to classify groups using support vector machine or predict individuals’ characteristics using support vector regression.

**Figure 2.**
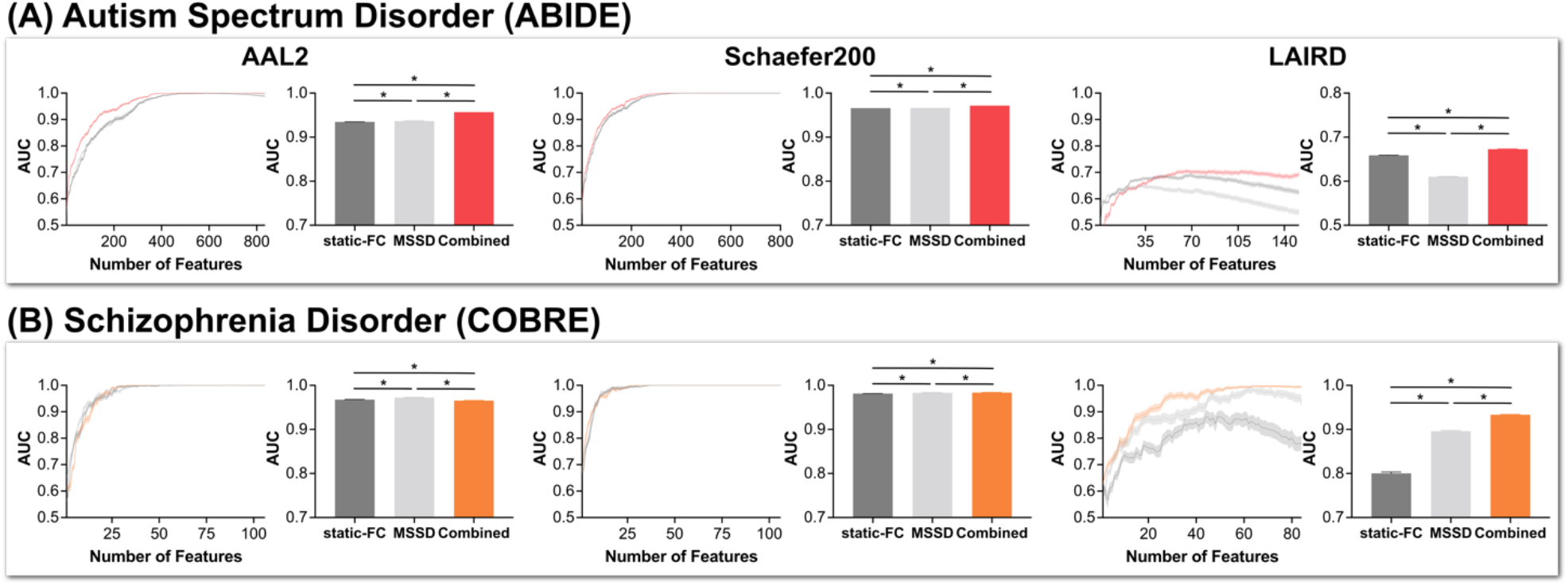
Group classification results for each pathological group using the support vector machine (SVM) with static-FC, MSSD, and the combined features. The classification performances were evaluated by the area under curve (AUC) first. Higher AUC means better classification performance. The line graphs on the left side represent overall classification performance as a function of the number of features when the smallest lambda was used in the feature selection stage. The bar graphs on the right side indicate the corresponding secondary AUC from the line graphs for statistical testing. The error bars are standard deviation (SD). (A) classification results of autism spectrum disorder and typical development. Regardless of brain parcellation, including FC-variability (i.e., combined) reached higher classification performances and the performances were more reliable. (B) classification results of schizophrenia disorder and the control. The ‘combined’ features showed slightly higher classification performances, but the amount of increased AUCs was small.

**Table 1.** Overall group classification performance using SVM

| Autism Spectrum Disorder (ABIDE) Classification |  |  |  |  |
| --- | --- | --- | --- | --- |
| Parcellation | static-FC | MSSD | combined | Permutation |
| AAL2 | 0.934 (± 0.0001) | 0.936 (± 0.0001) | 0.957 (± 0.0000) | $p < .001$ |
| Schaefer200 | 0.966 (± 0.0001) | 0.966 (± 0.0000) | 0.972 (± 0.0000) | $p < .001$ |
| LAIRD | 0.658 (± 0.0001) | 0.610 (± 0.0006) | 0.672 (± 0.0005) | $p < .001$ |
| Schizophrenia Disorder (COBRE) Classification |  |  |  |  |
| Parcellation | static-FC | MSSD | combined | Permutation |
| AAL2 | 0.967 (± 0.0005) | 0.972 (± 0.0004) | 0.965 (± 0.0005) | $p < .001$ |
| Schaefer200 | 0.981 (± 0.0004) | 0.983 (± 0.0003) | 0.984 (± 0.0003) | $p < .001$ |
| LAIRD | 0.800 (± 0.0028) | 0.896 (± 0.0016) | 0.933 (± 0.001) | $p < .001$ |

*Note:* Classification performances were evaluated by using the area under curve (AUC) first and then secondary AUCs from line graphs were calculated for statistical testing. Each cell represents the mean of secondary AUC ± SD. 'Permutation' means the permutation test results between the static-FC and the combined.

### Predictions of individuals characteristics

The SVR results also revealed that including the FC-variability was helpful to predict individuals’ characteristics. Across all SVR results, the ‘combined’ showed significantly lower secondary AUC than that of static-FC (all *p* < .001; See Figure 3 and Table 2). These results suggested that psychiatric symptoms are associated with FC-variability partially, which has been largely overlooked. Similar to the classification results, the predictive results were more reliable when FC-variability features were added. This was consistent when the optimal lambda was used (Supplementary Figure 1). The weights of SVR were presented in Figure 4 and Supplementary Figure 5-8.

**Figure 3.**
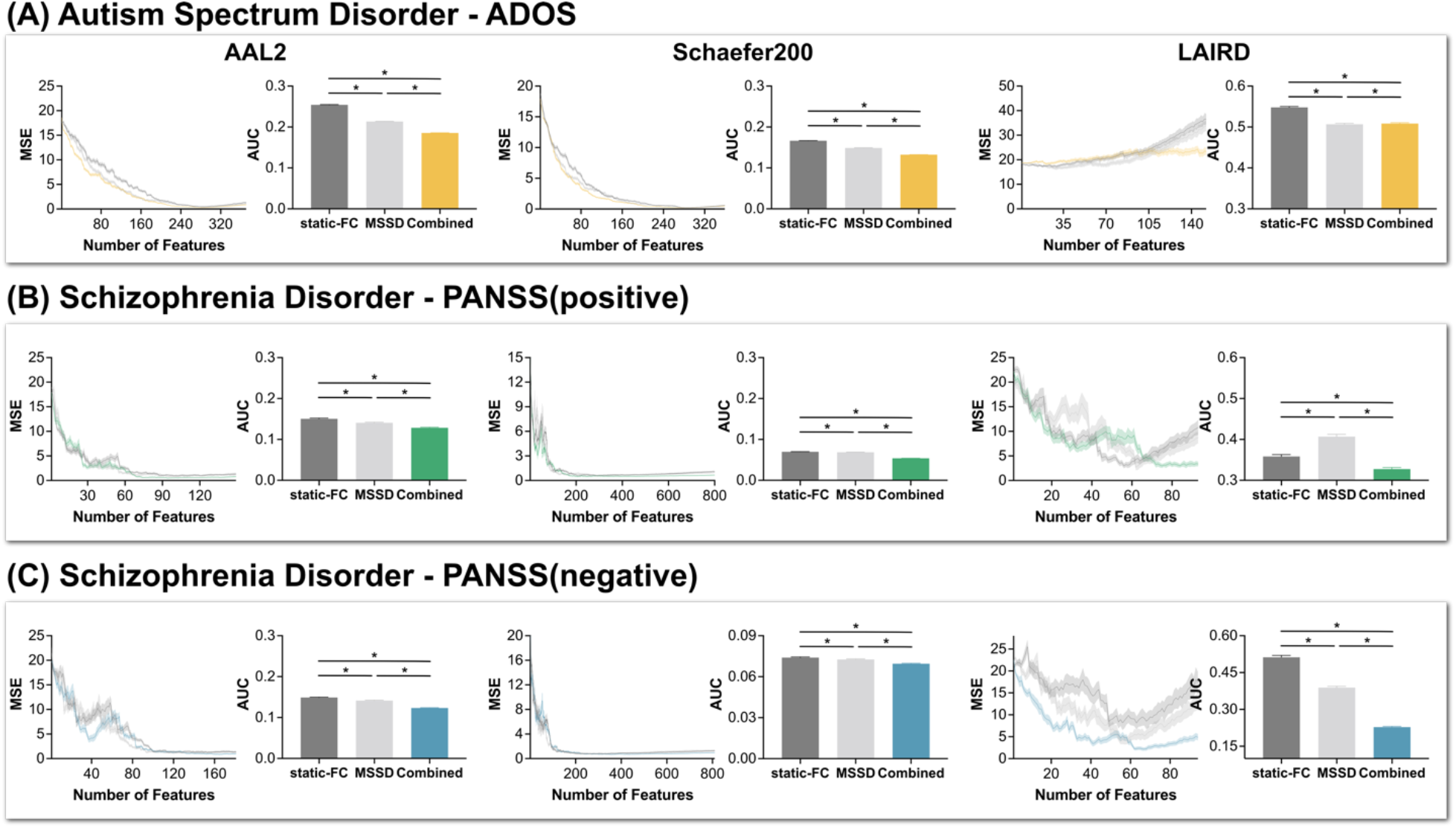
Prediction results using the support vector regression (SVR) with static-FC, MSSD, and the combined features. The predicted variables included (A) ADOS and (B) total scores of positive symptoms and (C) negative symptoms of PANSS. The prediction performances were evaluated by mean squared error (MSE) first. Lower MSE means better prediction. The line graphs on the left side represent overall prediction performance as a function of the number of features when the smallest lambda was used in the feature selection stage. In addition, the combined showed more reliable predictive results. The bar graphs on the right side indicate the corresponding secondary AUC from the line graphs for statistical testing. The error bars are standard deviation (SD).

**Figure 4.**
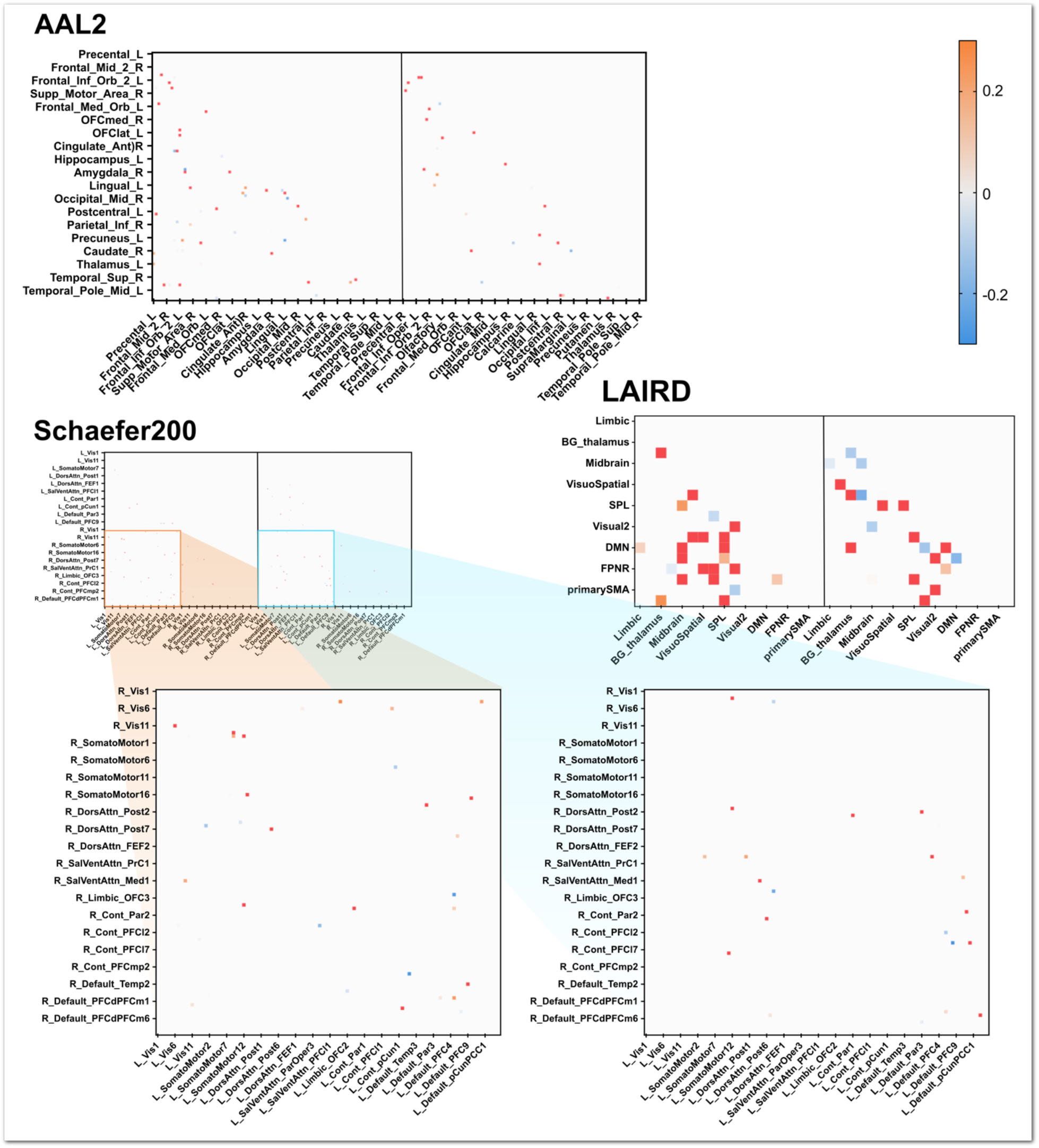
Weights of SVR which predicted the ADOS score. Notably, the weight patterns of FC-variability were different from the pattern of static-FC, suggesting that FC-variability has some independent and additional information. In each matrix, left square represents static-FC parts and right square represents FC-variability parts. More contributing features (weight > 0.3 or < −0.3) were colored red. Due to its enormous number of features, the weights of Schaefer200 atlas are presented here partially. Whole weight matrix is presented in Supplementary Figure 5.

**Table 2.** Overall predictive performance using SVR

| Autism Spectrum Disorder - ADOS |  |  |  |  |
| --- | --- | --- | --- | --- |
| Parcellation | Static-FC | MSSD | Combined | Permutation |
| AAL2 | 0.254 (± 0.0006) | 0.213 (± 0.0005) | 0.185 (± 0.0005) | $p < .001$ |
| Schaefer200 | 0.166 (± 0.0005) | 0.140 (± 0.0004) | 0.141 (± 0.0004) | $p < .001$ |
| LAIRD | 0.548 (±<br>0.0023) | 0.507 (±<br>0.0024) | 0.508 (±<br>0.0021) | $p < .001$ |
| Schizophrenia Disorder – PANSS(positive) |  |  |  |  |
| Parcellation | Static-FC | MSSD | Combined | Permutation |
| AAL2 | 0.150 (±<br>0.0016) | 0.141 (±<br>0.0015) | 0.128 (±<br>0.0015) | $p < .001$ |
| Schaefer200 | 0.069 (±<br>0.0003) | 0.068 (±<br>0.0004) | 0.054 (±<br>0.0003) | $p < .001$ |
| LAIRD | 0.358 (±<br>0.0045) | 0.407 (±<br>0.0055) | 0.327 (±<br>0.0039) | $p < .001$ |
| Schizophrenia Disorder – PANSS(negative) |  |  |  |  |
| Parcellation | Static-FC | MSSD | Combined | Permutation |
| AAL2 | 0.149 (±<br>0.0010) | 0.141 (±<br>0.0011) | 0.123 (±<br>0.0009) | $p < .001$ |
| Schaefer200 | 0.074 (±<br>0.0004) | 0.072 (±<br>0.0004) | 0.069 (±<br>0.0004) | $p < .001$ |
| LAIRD | 0.512 (±<br>0.0075) | 0.389 (±<br>0.0052) | 0.228 (±<br>0.0027) | $p < .001$ |

*Note:* Prediction performances were evaluated by using mean squared error (MSE) first and then secondary AUCs from line graphs were calculated for statistical testing. Lower secondary AUC means better SVR prediction performance. Each cell represents the mean of secondary AUC ± SD. 'Permutation' means the permutation test results between the static-FC and the combined.

### Identifying important FC-variability features

We investigated which FC-variability was important for the classifications and predictions. As a useful example, Figure 4 shows the weights of SVR which predicted the ADOS score with the optimal lambda. Other weights results are presented in the Supplementary Figure 3-8. Since the main aim of this analysis is to identify which FC-variability is informative when it was added to static-FC, we interpret the weights of the ‘combined’. In each weight matrix in Figure 4, the left square represents static-FC parts and the right square represents FC-variability parts. It should be noted again that these features were used simultaneously for the feature-selection and the subsequent ML (i.e., combined). More contributing features (weight > 0.3 or < −0.3) were colored red. Most importantly, the patterns of weights were different between static-FC and FC-variability, implying that FC-variability has independent and additional information than static-FC. In AAL2, many FC-variability between ROIs within frontal areas and orbitofrontal cortex (OFC) contributed to predict the ADOS score. Other brain regions were also involved such as amygdala, insula, caudate, postcentral gyrus, paracentral gyrus, inferior temporal areas, and occipital areas. Similarly, contributing FC-variability was dispersed across many lobes when Schaefer200 atlas was used. Interestingly, when we reviewed red-colored Schaefer features in the context of Yeo 7 networks which the developers had identified, the contributing FC-variability features were more derived from between-network ROIs (15/17) than the static-FC features (10/17). Although this pattern was not tested statistically, the LAIRD network atlas result supported the pattern by showing that numerous between-network FC-variability was important to predict the ADOS score.

## Discussion

To understand brain’s functional system, the static-FC has been widely used to estimate FC between brain regions but resulted in not reflecting connectivity fluctuations. The Dynamic-FC, thus, has been suggested as a promising method to capture FC fluctuation across time and has proved that the brain has several mental states even when external stimuli are absent. However, much fewer studies utilized the FC-variability and few studies compared the static-FC and the FC-variability (Fong et al., 2019; Wang et al., 2018). This study aimed to demonstrate the benefits of FC-variability and to prove that the benefits are consistent across various cognitive domains and analytic procedures.

First, the current study classified the ASD and the TD using the ABIDE and classified the schizophrenia and the control using the COBRE. Regardless of parcellation selection, including the FC-variability increased classification performance mostly (Figure 2, Supplementary Figure 1). In addition, FC-variability was also beneficial to predict individuals’ characteristics including ADOS, PANSS, and age. The beneficial effects were robust across various analytic procedures again. In sum, these results suggested that FC-variability may be relevant to various cognitive domains and can be informative across various analytic procedures. Second, we also identified which components of FC-variability were informative for classifying and predicting (Figure 4 and Supplementary Figure 3-8). In general, a similar number of features of FC-variability compared to the static-FC showed a high contribution to classification and prediction. Furthermore, It is also noteworthy that the contributing FC-variability was in the different locations implying that the FC-variability has independent information in comparison to the static-FC.

Compared to the previous findings that utilized FC-variability, the current findings of classification were somewhat different. For instance, Y. Li et al. (2020) reported that the FC-variability of inferior frontal gyrus (opecular part) - post cingulate cortex was significantly different between the ASD and the TD; that FC-variability was not replicated in the current study, we observed several significant OFC-related FC-variability when we reviewed the ‘combined’ feature (Supplementary Fig 3). Similarly, for the schizophrenia classification, Supekar et al. (2019) suggested a triple-network saliency model that emphasized dynamic cross-network interactions between salience network, central executive network, and default mode network. In the current result, FC-variability of Rectus – Amygdala and anterior OFC – Occipital area contributed to classify the groups (Supplementary Figure 4). Due to the absence of research associating FC-variability and psychiatric scores including ADOS and PANSS, however, these results could not be compared with previous findings. We speculate that the main reason of discrepancy was that we used both static-FC and FC-variability and then FC-variability features which had redundant information were less noticeable. Nevertheless, we believe that including more independent and informative features is crucial to utilize neuroimages in psychiatric field such as connectome-based predictive modeling (CPM; Liu et al., 2022; Tian & Zalesky, 2021). Another possibility of discrepancy was that we adopted robust correlation method to estimate static-FC and FC-variability. In our empirical comparisons, the robust method significantly increased static-FC performance (Supplementary Fig 9).

Recent studies imply that different gene expressions can be a factor that drives the different degree of FC-variability across individuals (Barber et al., 2021; Liu et al., 2019). Specifically, Liu et al. (2019) estimated FC-variability brain maps from rs-fMRI of Human Connectome Project and UK Biobank, compared them with gene expression profiles from Allen Human Brain Atlas^3^, and found that FC-variability had a significant relationship with two groups of gene expressions: one was synaptic plasticity processes such as action potential, ion transportation and hormone secretion. Such gene expressions affect synaptic transmission processes especially at the molecular level and are related to fast FC changes (Arnsten et al., 2010). Another was structural plasticity such as the formation, development, and reorganization of presynaptic and postsynaptic constructs. These processes also influence synaptic plasticity and subsequent FC, such as cGMP-mediated signaling regulates synaptic plasticity. (Kleppisch & Feil, 2009). Such neuroplasticity has been linked to psychiatric disorders including ASD and schizophrenia (Bernardinelli et al., 2014), our conjecture is that the current informative FC-variability features reflected several genetic alterations of psychiatric disorders.

Finally, we would like to note some suggestions which should be addressed in the future studies. The current study demonstrated the robustness of FC-variability benefits regardless of time window sizes of sliding-window approach which is most widely used. However, there are several recently developed methods to calculate dynamic-FC such as dynamic conditional correlation (DCC; Choe et al., 2017; Lindquist et al., 2014) and edge-centric time series (Faskowitz et al., 2020; Zamani Esfahlani et al., 2020), and such methods should be compared in the context of FC-variability. Furthermore, we used three widely used atlases including structure-based (AAL2), functional-based (Schaefer), and network-based (LAIRD) atlases. Our results demonstrated that FC-variability is beneficial regardless of atlas selection, but there is a possibility that the benefits can be increased when the brain is parcellated into regions showing homogeneous FC-variability degree. Most recently, a few studies developed brain atlas considering FC-variability (Fan et al., 2021; Peng et al., 2022), it would be worthwhile to compare it with the conventional atlases.

We demonstrated the benefits of FC-variability across many cognitive domains, analytic variants, and datasets. Adding FC-variability not only increased ML performances also resulted in more reliable results. We suggest that many psychological and transdiagnostic research field such as connectome-based predictive modeling utilize FC-variability and hope that the findings may shed light on neuroimaging field.

## Supplementary Document

**Supplementary Figure 1.**
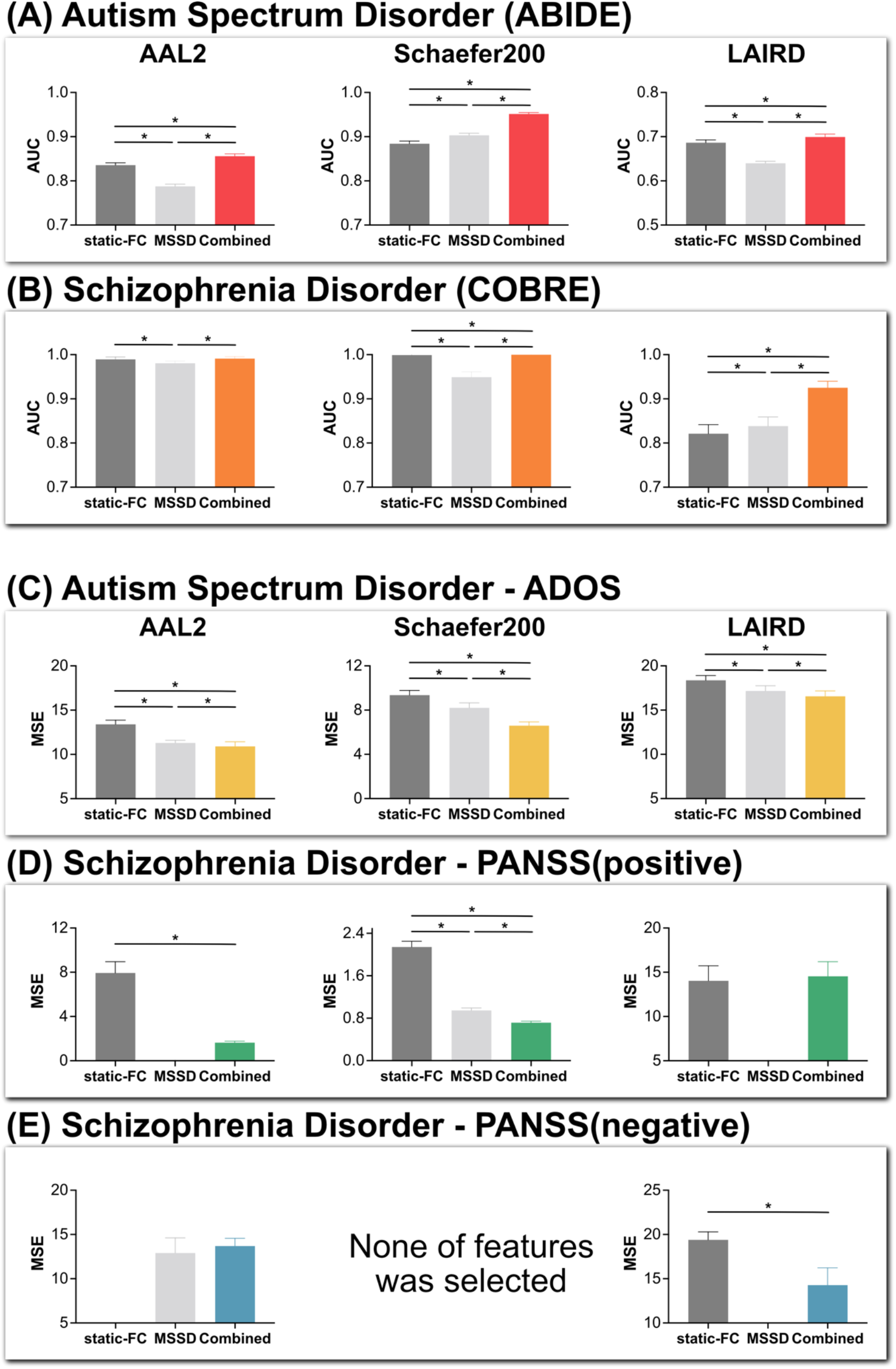
ML results using the optimal lambda at the feature selection stage. (A-B): Group classification results using SVM with static-FC, MSSD, and the combined features. The classification performances were evaluated by AUC. Higher AUC means better classification performance. The results show that including FC-variability (i.e., combined) resulted in higher classification performance. (C-E): Predictive results using SVR. Performances were evaluated by MSE. Lower MSE indicates better prediction. Null results mean no feature was selected at the feature selection stage. Again, including FC-variability increased predictive performance. The error bars are standard deviation (SD).

**Supplementary Figure 2.**
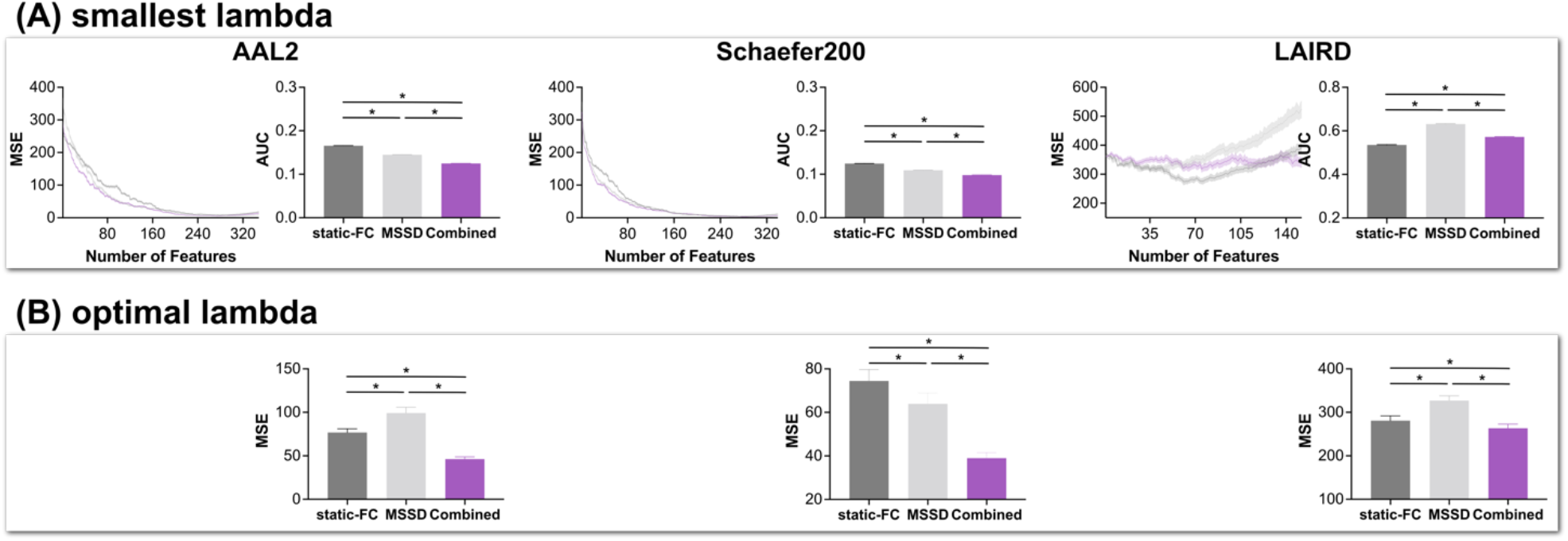
Individuals’ age prediction using NKI dataset and SVR. Lower values mean better predictive performance. (A) predictive performance with smallest lambda. The bar graphs on the right side show the corresponding secondary AUC of the line graphs. (B) predictive performance with the optimal lambda. Mostly, including FC-variability was beneficial to predict age. One exception was the overall predictive performance (i.e., secondary AUC) when the LAIRD atlas was used due to better performances of static-FC at some points. However, including FC-variability still showed more reliable predictive performance compared to the static-FC solely.

**Supplementary Figure 3.**
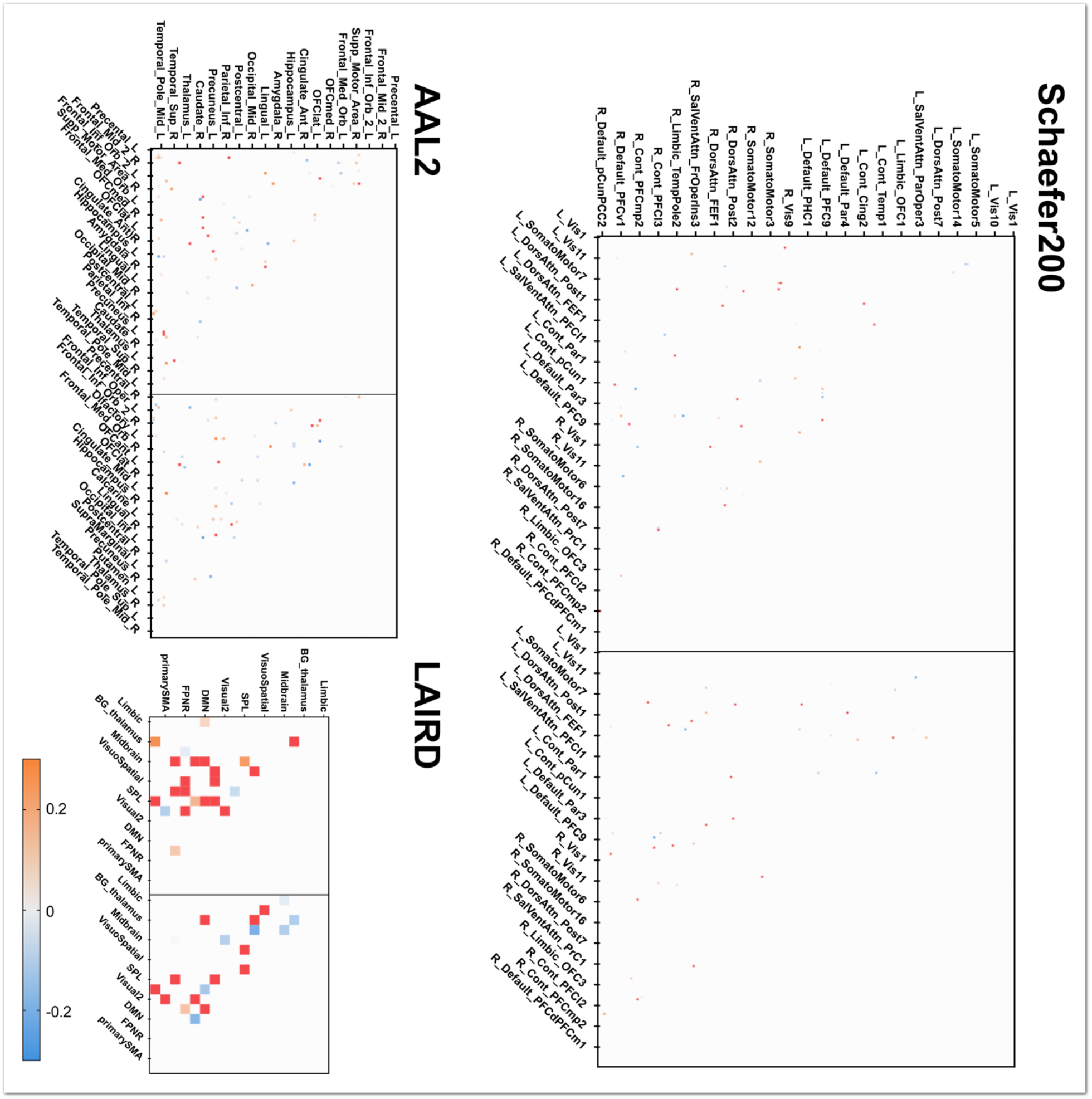
Weights of SVM which classified the autistic participants and the typical development (TD). In each matrix, left square represents of static-FC parts and right square represents FC-variability parts. More contributed features (weight > 0.3 or weight < −0.3) are colored red. The most contributing FC-variability features of AAL2 were Frontal_Inf_Orb - OFCpost, Rolandic_Oper - Insula, OFCant - Angular, OFCpost - Heschl, Lingual - Parietal_Sup, and Occipital_Mid - Angular. In the Schaefer200 result, many contributing features were associated with ROIs within somatomotor, control, default mode networks. In LAIRD network level, FC-variability of SMA – Visual3, Visuospatial – Visual 3 was informative to classify the groups.

**Supplementary Figure 4.**
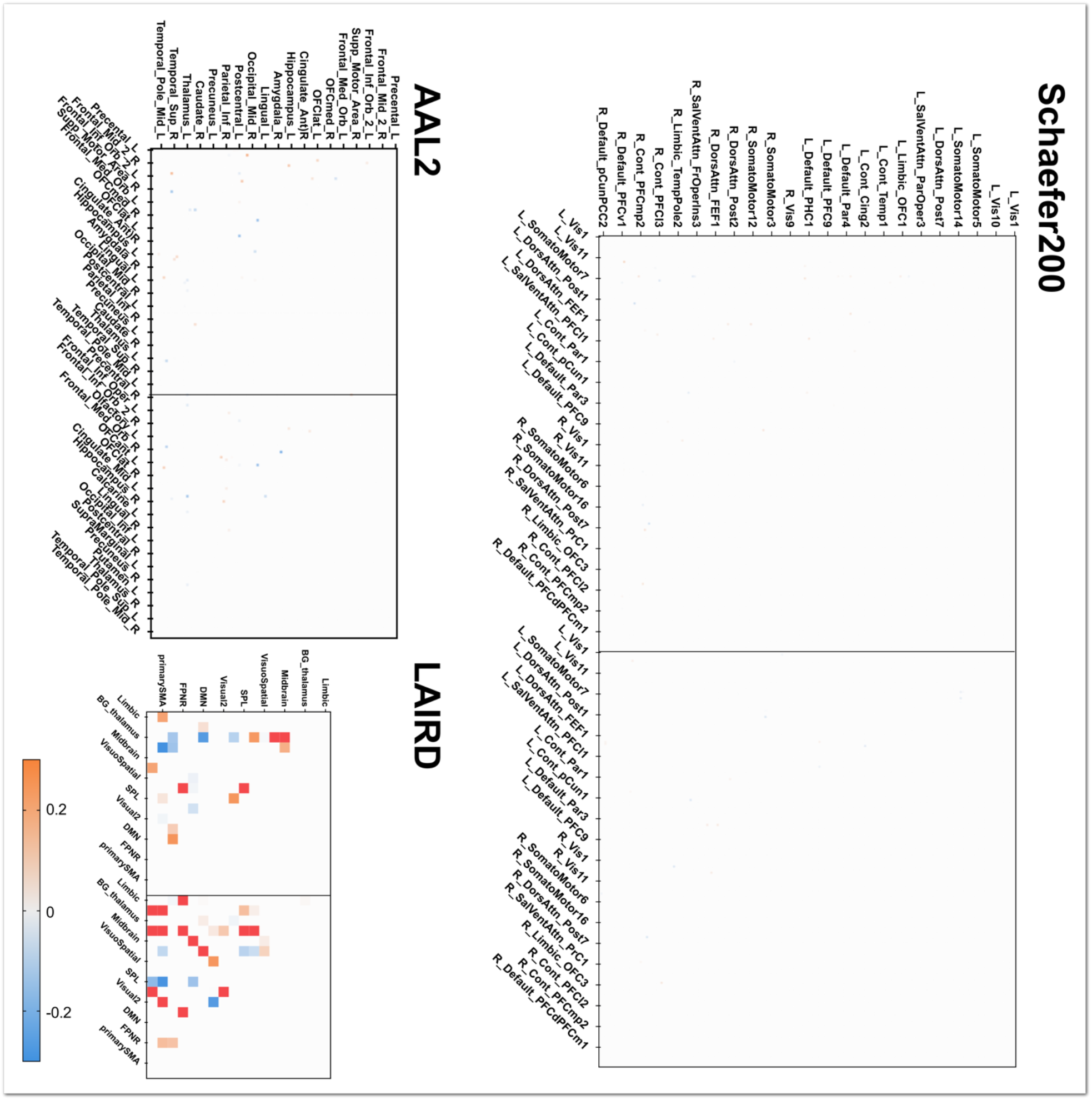
Weights of SVM to classify the schizophrenia and the control. In each matrix, left square represents of static-FC parts and right square represents FC-variability parts. More contributed features (weight > 0.3 or weight < −0.3) are colored red. Compared to other classification and predictions, less features contributed to classify the schizophrenia and the control. Nonetheless, some pairs contributed to the classification such as Rectus – Amygdala, OFCant – Occipital_Sup (AAL2). In LAIRD network level, some FC-variability showed high contributions additionally; Limbic – Visuospatial, SMA – Visual3, and Visuospatial – Visual3. In Schaefer200, any feature was not prominent to classify the groups.

**Supplementary Figure 5.**
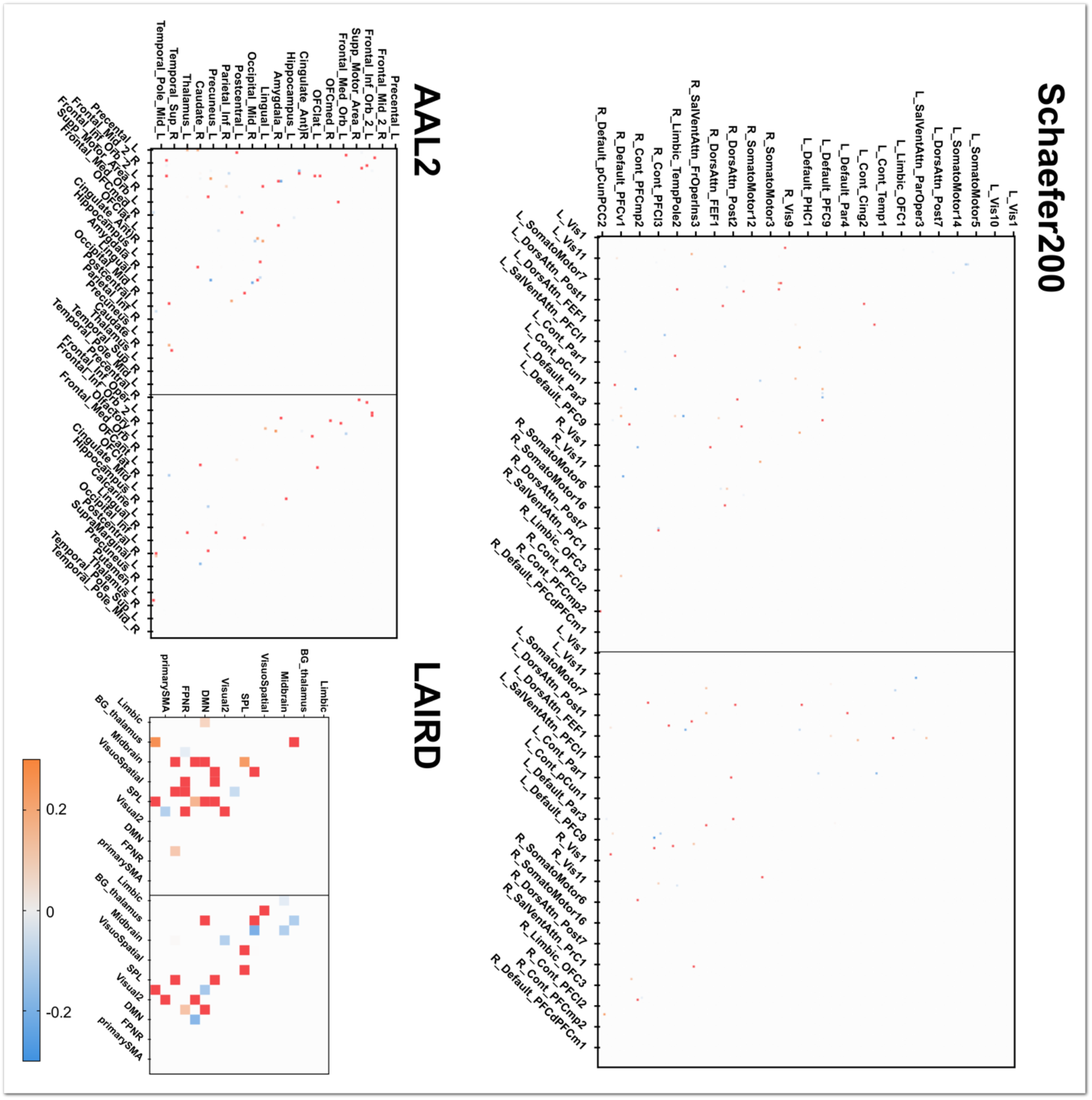
Weights of SVR to predict ADOS scores of autistic individuals. In each matrix, left square represents of static-FC parts and right square represents FC-variability parts. More contributed features (weight > 0.3 or weight < −0.3) are colored red. Many FC-variability between ROIs within frontal areas and orbitofrontal cortex (OFC) contributed to predict the ADOS score. Other brain regions were also involved such as amygdala, insula, caudate, postcentral gyrus, paracentral gyrus, inferior temporal areas, and occipital areas.

**Supplementary Figure 6.**
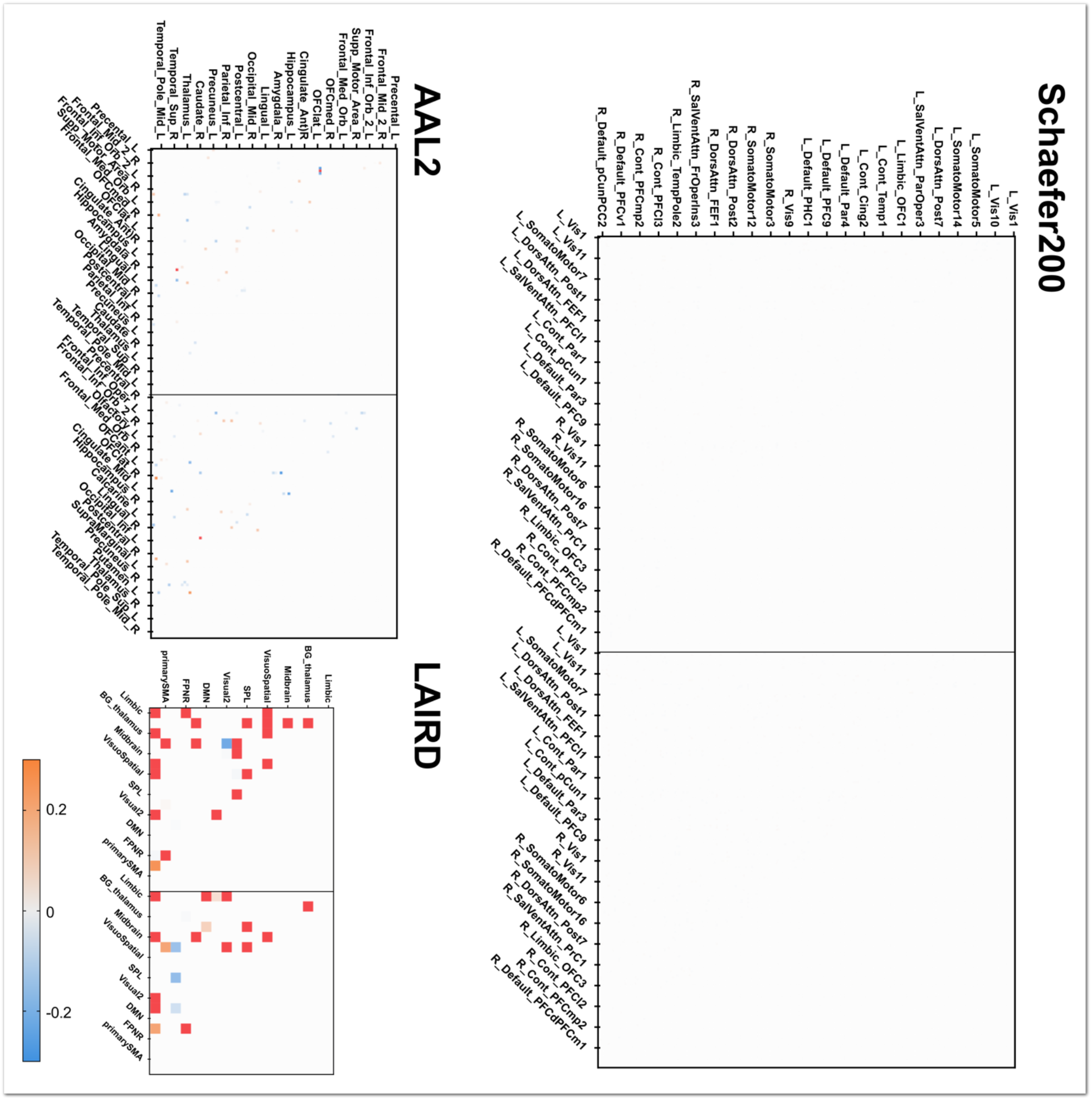
Weights of SVR to predict PANSS (positive total) scores of schizophrenic individuals. In each matrix, left square represents of static-FC parts and right square represents FC-variability parts. More contributed features (weight > 0.3 or weight < −0.3) are colored red. Importantly, several FC-variabililty were associated with the positive symptoms: Occipital_Mid – Caudate, OFClat – Amygdala, OFCant – Pallidum, Cingulate_Post – Hippocampus (AAL2). In the network level, limbic network and OFC-related network showed high contributions.

**Supplementary Figure 7.**
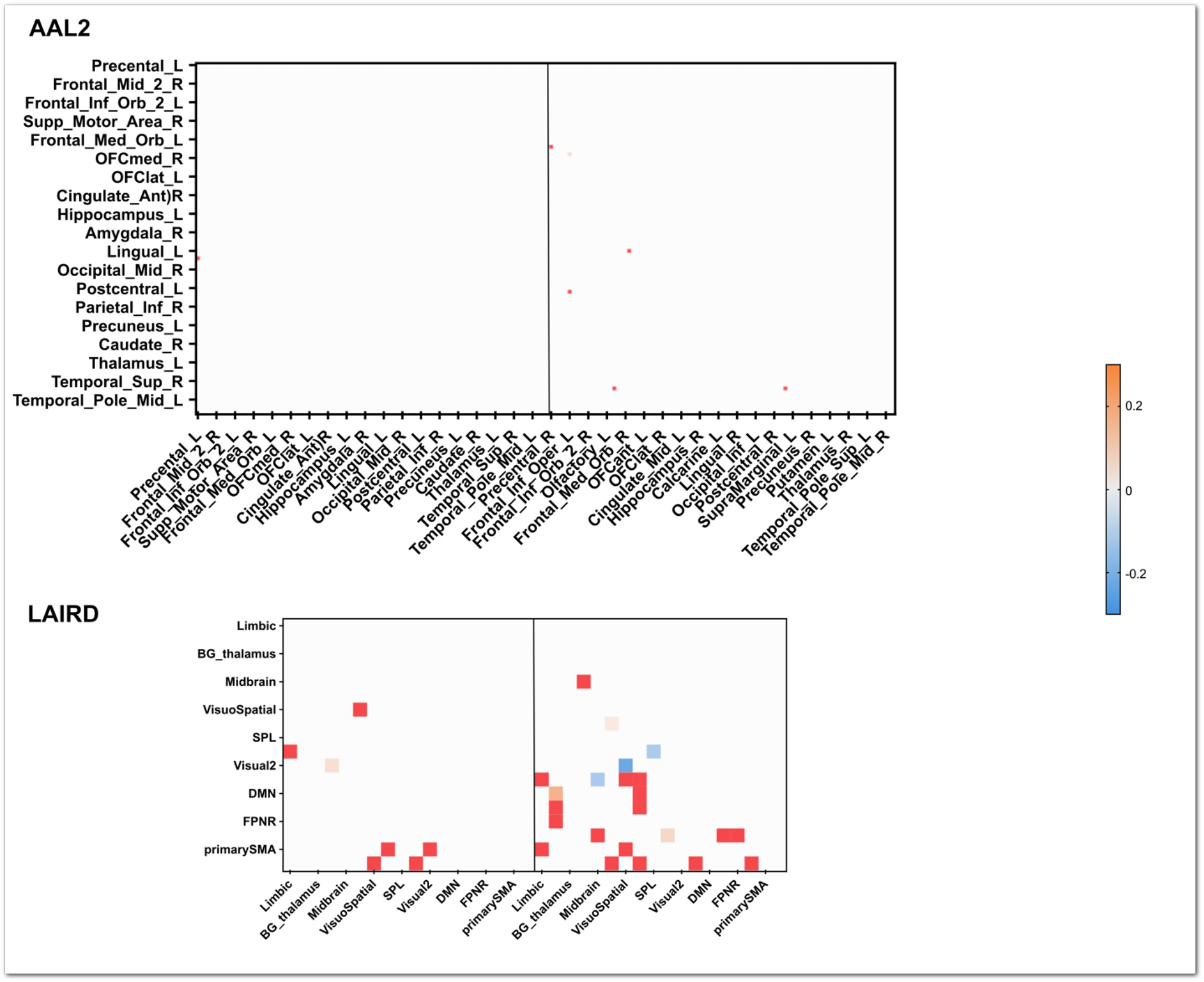
Weights of SVR to predict PANSS (negative total) scores of schizophrenic individuals. In each matrix, left square represents of static-FC parts and right square represents FC-variability parts. More contributed features (weight > 0.3 or weight < −0.3) are colored red. No features were selected when the optimal lambda and Schaefer200 atlas were used. Interestingly, in general, the negative symptoms were more associated with the FC-variability than the static-FC. In AAL2, Precentral – Rectus, Rectus – Lingual, Frontal_Inf_Oper – Pareital_Sup, Frontal_Sup_Medial – Temporal_Pole, and Parietal_Inf – Temporal_Pole was involved. In the LAIRD, limbic, OFC, and motor networks showed high contributions to predict the negative symptoms.

**Supplementary Figure 8.**
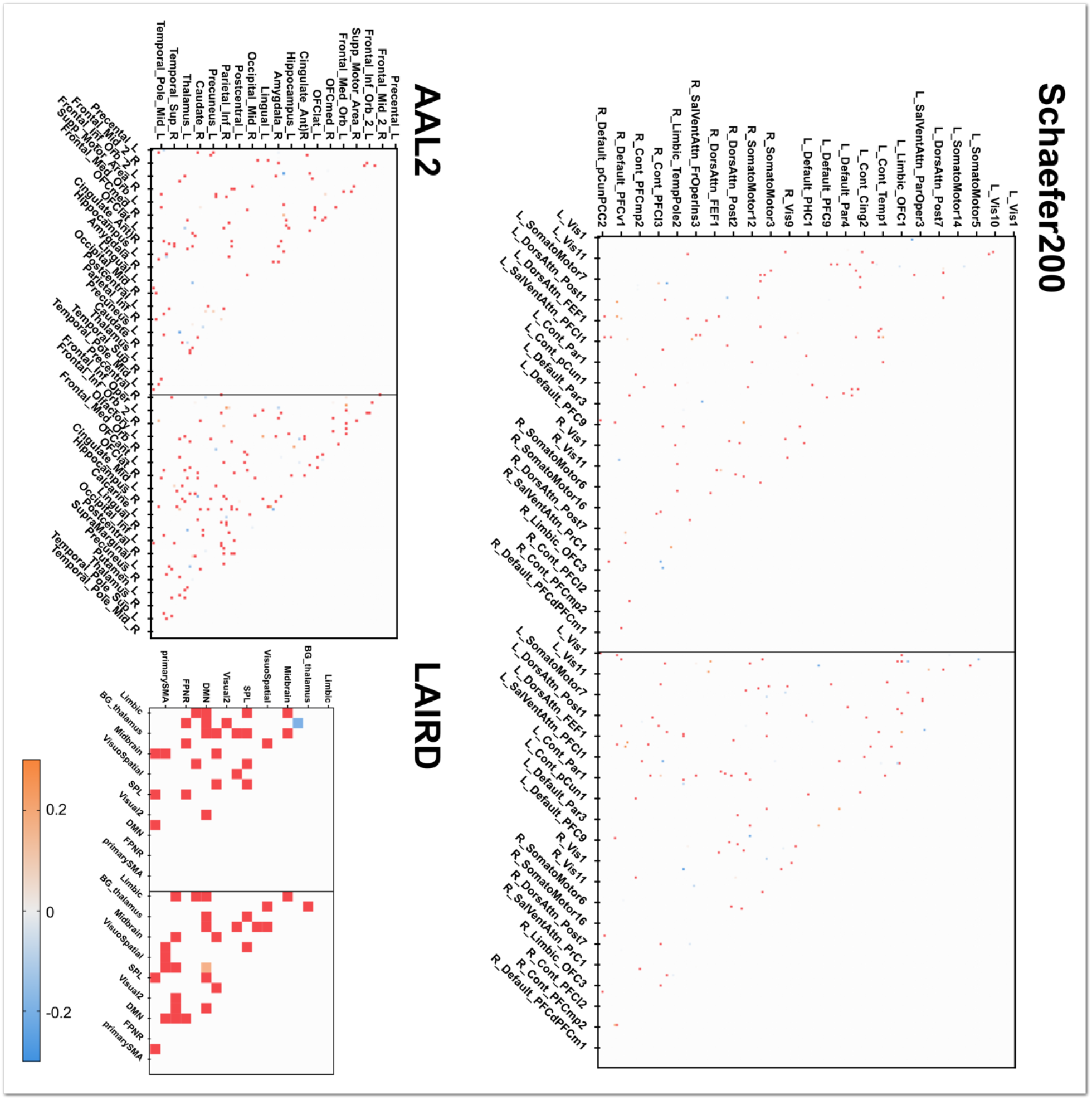
Weights of SVR to predict individuals’ age from the NKI dataset. In each matrix, left square represents of static-FC parts and right square represents FC-variability parts. More contributed features (weight > 0.3 or weight < −0.3) are colored red. Compared to the previous three psychiatric symptom predictions, much more FC-variability features were related to the individuals’ age, suggesting that the ageing process is not limited to the changes of some brain regions. Importantly, most of contributing FC-variability features to predict age did not overlap with the contributing static-FC features.

**Supplementary Figure 9.**
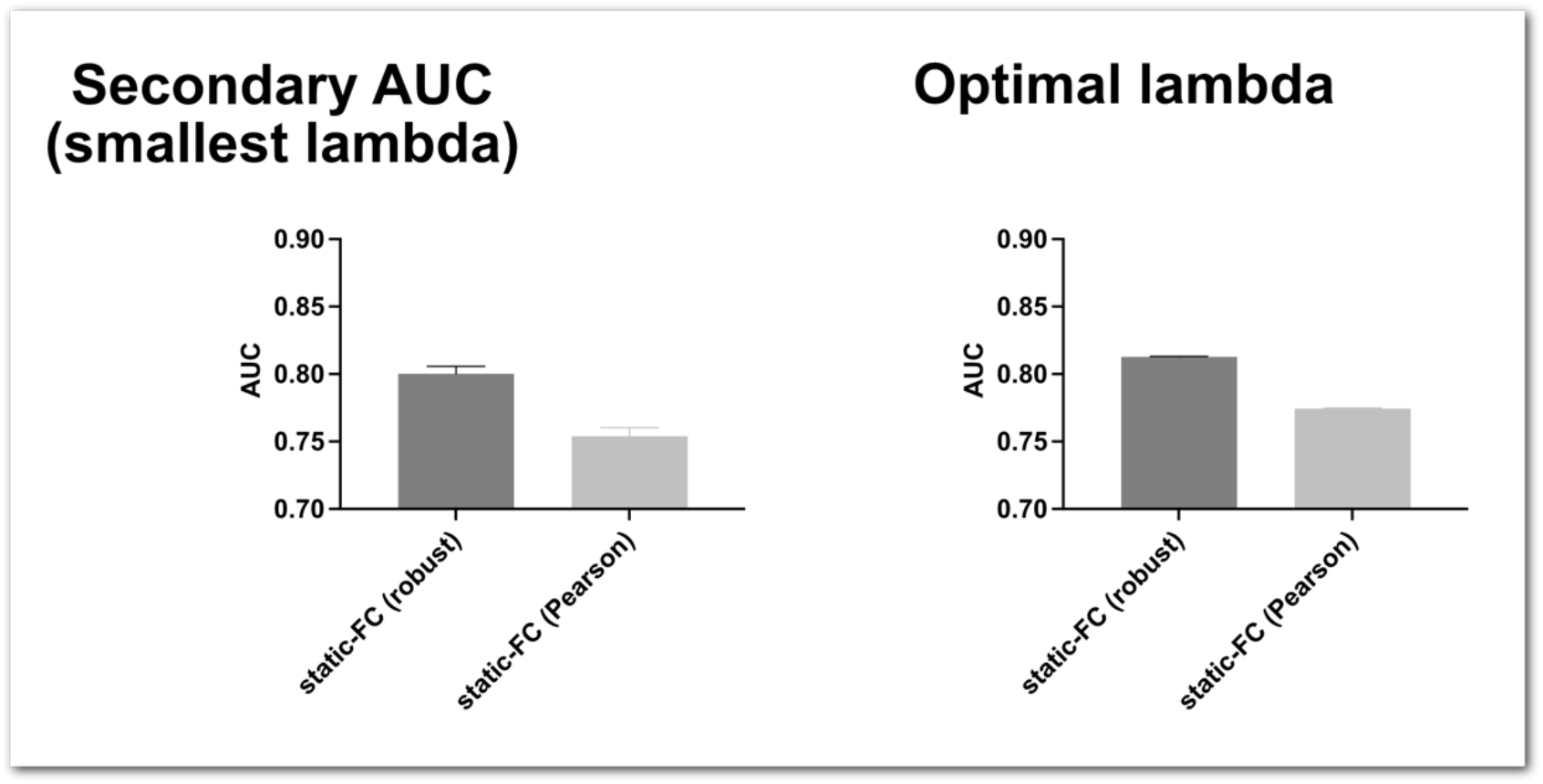
Comparison of robust correlation and Pearson correlation when classifying ABIDE (autism vs typical development) dataset with static-FC solely. The robust correlation method showed significantly higher classification performance than Pearson correlation.

**Supplementary Table 1.** Group classification performance with various sliding-window sizes (smallest lambda)

| Autism Spectrum Disorder (ABIDE) Classification |  |  |  |  |
| --- | --- | --- | --- | --- |
| Sliding-window size: 60s |  |  |  |  |
| Parcellation | static-FC | MSSD | combined | Permutation |
| AAL2 | 0.934 (± 0.0001) | 0.936 (± 0.0001) | 0.957 (± 0.0000) | $p < .001$ |
| Schaefer200 | 0.966 (± 0.0001) | 0.966 (± 0.0000) | 0.972 (± 0.0000) | $p < .001$ |
| LAIRD | 0.658 (± 0.0001) | 0.610 (± 0.0006) | 0.672 (± 0.0005) | $p < .001$ |
| Sliding-window size: 90s |  |  |  |  |
| AAL2 | 0.934 (± 0.0001) | 0.925 (± 0.0001) | 0.951 (± 0.0001) | $p < .001$ |
| Schaefer200 | 0.966 (± 0.0001) | 0.961 (± 0.0001) | 0.971 (± 0.0001) | $p < .001$ |
| LAIRD | 0.658 (± 0.0007) | 0.610 (± 0.0007) | 0.685 (± 0.0006) | $p < .001$ |
| Sliding-window size: 120s |  |  |  |  |
| AAL2 | 0.934 (± 0.0001) | 0.925 (± 0.0001) | 0.951 (± 0.0001) | $p < .001$ |
| Schaefer200 | 0.966 (± 0.0001) | 0.965 (± 0.0001) | 0.970 (± 0.0000) | $p < .001$ |
| LAIRD | 0.658 (± 0.0007) | 0.604 (± 0.0007) | 0.678 (± 0.0006) | $p < .001$ |
| Schizophrenia Disorder (COBRE) Classification |  |  |  |  |
| Sliding-window size: 60s |  |  |  |  |
| Parcellation | static-FC | MSSD | combined | Permutation |
| AAL2 | 0.967 (± 0.0005) | 0.972 (± 0.0004) | 0.965 (± 0.0005) | $p < .001$ |
| Schaefer200 | 0.981 (± 0.0004) | 0.983 (± 0.0003) | 0.984 (± 0.0003) | $p < .001$ |
| LAIRD | 0.800 (± 0.0028) | 0.896 (± 0.0016) | 0.933 (± 0.001) | $p < .001$ |
| Sliding-window size: 90s |  |  |  |  |
| AAL2 | 0.967 (± 0.0005) | 0.977 (± 0.0004) | 0.969 (± 0.0005) | $p < .001$ |
| Schaefer200 | 0.981 (± 0.0003) | 0.977 (± 0.0004) | 0.983 (± 0.0003) | $p < .001$ |
| LAIRD | 0.801 (± 0.0021) | 0.855 (± 0.0021) | 0.909 (± 0.0012) | $p < .001$ |
| Sliding-window size: 120s |  |  |  |  |
| AAL2 | 0.967 (± 0.0004) | 0.974 (± 0.0004) | 0.975 (± 0.0004) | $p < .001$ |
| Schaefer200 | 0.981 (± 0.0004) | 0.985 (± 0.0003) | 0.980 (± 0.0005) | $p < .001$ |
| LAIRD | 0.800 (± 0.0027) | 0.846 (± 0.0024) | 0.890 (± 0.0015) | $p < .001$ |

*Note:* Classification performances were evaluated by using the area under curve (AUC) first and the secondary AUC was calculated to evaluate overall performance. Each cell represents the mean of secondary AUC ± SD. Higher values indicate better overall performance. 'Permutation' means the permutation test results between the static-FC and the combined.

**Supplementary Table 2.** Predicting individuals’ characteristics performance with various sliding-window sizes (smallest lambda)

| Autism Spectrum Disorder – ADOS |  |  |  |  |
| --- | --- | --- | --- | --- |
| Sliding-window size: 60s |  |  |  |  |
| Parcellation | static-FC | MSSD | combined | Permutation |
| AAL2 | 0.254 (±<br>0.0006) | 0.213 (±<br>0.0005) | 0.185 (±<br>0.0005) | $p < .001$ |
| Schaefer200 | 0.166 (±<br>0.0005) | 0.140 (±<br>0.0004) | 0.141 (±<br>0.0004) | $p < .001$ |
| LAIRD | 0.548 (±<br>0.0023) | 0.507 (±<br>0.0024) | 0.508 (±<br>0.0021) | $p < .001$ |
| Sliding-window size: 90s |  |  |  |  |
| AAL2 | 0.260 (±<br>0.0007) | 0.203 (±<br>0.0005) | 0.203 (±<br>0.0005) | $p < .001$ |
| Schaefer200 | 0.167 (±<br>0.0005) | 0.140 (±<br>0.0003) | 0.138 (±<br>0.0006) | $p < .001$ |
| LAIRD | 0.517 (±<br>0.0022) | 0.501 (±<br>0.0021) | 0.501 (±<br>0.0018) | $p < .001$ |
| Sliding-window size: 120s |  |  |  |  |
| AAL2 | 0.261 (±<br>0.0008) | 0.203 (±<br>0.0005) | 0.186 (±<br>0.0004) | $p < .001$ |
| Schaefer200 | 0.164 (±<br>0.0004) | 0.149 (±<br>0.0003) | 0.148 (±<br>0.0003) | $p < .001$ |
| LAIRD | 0.409 (±<br>0.0013) | 0.472 (±<br>0.0020) | 0.448 (±<br>0.0018) | $p < .001$ |
| Schizophrenia Disorder – PANSS(positive) |  |  |  |  |
| Sliding-window size: 60s |  |  |  |  |

| Parcellation | static-FC | MSSD | Combined | Permutation |
| --- | --- | --- | --- | --- |
| AAL2 | 0.150 (±<br>0.0016) | 0.141 (±<br>0.0015) | 0.128 (±<br>0.0015) | $p < .001$ |
| Schaefer200 | 0.069 (±<br>0.0003) | 0.068 (±<br>0.0004) | 0.054 (±<br>0.0003) | $p < .001$ |
| LAIRD | 0.358 (±<br>0.0045) | 0.407 (±<br>0.0055) | 0.327 (±<br>0.0039) | $p < .001$ |
| Sliding-window size: 90s |  |  |  |  |
| AAL2 | 0.158 (±<br>0.0016) | 0.146 (±<br>0.0015) | 0.137 (±<br>0.0015) | $p < .001$ |
| Schaefer200 | 0.067 (±<br>0.0003) | 0.070 (±<br>0.0005) | 0.062 (±<br>0.0004) | $p < .001$ |
| LAIRD | 0.360 (±<br>0.0042) | 0.357 (±<br>0.0044) | 0.285 (±<br>0.0033) | $p < .001$ |
| Sliding-window size: 120s |  |  |  |  |
| AAL2 | 0.146 (±<br>0.0012) | 0.164 (±<br>0.0022) | 0.118 (±<br>0.0012) | $p < .001$ |
| Schaefer200 | 0.065 (±<br>0.0003) | 0.080 (±<br>0.0005) | 0.057 (±<br>0.0002) | $p = 0.186$ |
| LAIRD | 0.319 (±<br>0.0039) | 0.399 (±<br>0.0048) | 0.289 (±<br>0.0035) | $p < .001$ |
| Schizophrenia Disorder – PANSS(negative) |  |  |  |  |
| Sliding-window size: 60s |  |  |  |  |
| Parcellation | static-FC | MSSD | Combined | Permutation |
| AAL2 | 0.149 (±<br>0.0010) | 0.141 (±<br>0.0011) | 0.123 (±<br>0.0009) | $p < .001$ |
| Schaefer200 | 0.074 (±<br>0.0004) | 0.072 (±<br>0.0004) | 0.069 (±<br>0.0004) | $p < .001$ |
| LAIRD | 0.512 (±<br>0.0075) | 0.389 (±<br>0.0052) | 0.228 (±<br>0.0027) | $p < .001$ |
| Sliding-window size: 90s |  |  |  |  |
| AAL2 | 0.170 (±<br>0.0021) | 0.128 (±<br>0.0015) | 0.140 (±<br>0.0016) | $p < .001$ |
| Schaefer200 | 0.075 (±<br>0.0005) | 0.069 (±<br>0.0004) | 0.069 (±<br>0.0005) | $p < .001$ |
| LAIRD | 0.460 (±<br>0.0054) | 0.302 (±<br>0.0035) | 0.296 (±<br>0.0039) | $p < .001$ |
| Sliding-window size: 120s |  |  |  |  |
| AAL2 | 0.168 (±<br>0.0019) | 0.177 (±<br>0.0019) | 0.152 (±<br>0.0016) | $p < .001$ |
| Schaefer200 | 0.076 (±<br>0.0004) | 0.083 (±<br>0.0004) | 0.073 (±<br>0.0004) | $p < .001$ |
| LAIRD | 0.466 (±<br>0.0056) | 0.351 (±<br>0.0052) | 0.231 (±<br>0.0028) | $p < .001$ |
| Neurodevelopmental (NKI)- Age |  |  |  |  |
| Sliding-window size: 60s |  |  |  |  |
| Parcellation | static-FC | MSSD | combined | Permutation |
| AAL2 | 0.166 (±<br>0.0003) | 0.145 (±<br>0.0004) | 0.125 (±<br>0.0002) | $p < .001$ |
| Schaefer200 | 0.125 (±<br>0.0002) | 0.109 (±<br>0.0003) | 0.098 (±<br>0.0002) | $p < .001$ |
| LAIRD | 0.535 (±<br>0.0016) | 0.631 (±<br>0.0018) | 0.572 (±<br>0.0018) | $p < .001$ |
| Sliding-window size: 90s |  |  |  |  |
| AAL2 | 0.164 (±<br>0.0004) | 0.168 (±<br>0.0004) | 0.130 (±<br>0.0002) | $p < .001$ |
| Schaefer200 | 0.130 (±<br>0.0003) | 0.127 (±<br>0.0003) | 0.110 (±<br>0.0002) | $p < .001$ |
| LAIRD | 0.529 (±<br>0.0014) | 0.625 (±<br>0.0019) | 0.573 (±<br>0.0014) | $p < .001$ |
| Sliding-window size: 120s |  |  |  |  |
| AAL2 | 0.165 (±<br>0.0003) | 0.189 (±<br>0.0005) | 0.132 (±<br>0.0003) | $p < .001$ |
| Schaefer200 | 0.125 (±<br>0.0002) | 0.128 (±<br>0.0003) | 0.113 (±<br>0.0002) | $p < .001$ |
| LAIRD | 0.526 (±<br>0.0019) | 0.624 (±<br>0.0022) | 0.544 (±<br>0.0015) | $p < .001$ |

*Note:* Prediction performances were evaluated by using mean squared error (MSE) first and the secondary AUC was calculated to evaluate overall performance. Lower secondary AUC means better SVR prediction performance. Each cell represents the mean of secondary AUC ± SD. 'Permutation' means the permutation test results between the static-FC and the combined.

**Supplementary Table 3.** Group classification performance with various sliding-window sizes (optimal lambda)

| Autism Spectrum Disorder (ABIDE) Classification |  |  |  |  |
| --- | --- | --- | --- | --- |
| Sliding-window size: 60s |  |  |  |  |
| Parcellation | static-FC | MSSD | combined | Permutation |
| AAL2 | 0.837 (± 0.006) | 0.789 (± 0.006) | 0.859 (± 0.004) | $p < .001$ |
| Schaefer200 | 0.884 (± 0.006) | 0.903 (± 0.005) | 0.951 (± 0.003) | $p < .001$ |
| LAIRD | 0.688 (± 0.007) | 0.638 (± 0.007) | 0.697 (± 0.008) | $p < .001$ |
| Sliding-window size: 90s |  |  |  |  |
| AAL2 | 0.837 (± 0.006) | 0.791 (± 0.003) | 0.871 (± 0.004) | $p < .001$ |
| Schaefer200 | 0.878 (± 0.007) | 0.871 (± 0.005) | 0.922 (± 0.005) | $p < .001$ |
| LAIRD | 0.678 (± 0.006) | 0.641 (± 0.007) | 0.716 (± 0.006) | $p < .001$ |
| Sliding-window size: 90s |  |  |  |  |
| AAL2 | 0.838 (± 0.006) | 0.789 (± 0.003) | 0.873 (± 0.007) | $p < .001$ |
| Schaefer200 | 0.878 (± 0.007) | 0.881 (± 0.005) | 0.919 (± 0.004) | $p < .001$ |
| LAIRD | 0.683 (± 0.007) | 0.622 (± 0.007) | 0.708 (± 0.006) | $p < .001$ |
| Schizophrenia Disorder (COBRE) Classification |  |  |  |  |
| Sliding-window size: 60s |  |  |  |  |
| Parcellation | static-FC | MSSD | combined | Permutation |
| AAL2 | 0.989 (± 0.004) | 0.979 (± 0.004) | 0.991 (± 0.004) | $p = .426$ |
| Schaefer200 | 0.999 (± 0.001) | 0.949 (± 0.012) | 0.999 (± 0.000) | $p < .001$ |
| LAIRD | 0.811 (± 0.019) | 0.847 (± 0.019) | 0.928 (± 0.013) | $p < .001$ |
| Sliding-window size: 90s |  |  |  |  |
| AAL2 | 0.986 (± 0.006) | 0.999 (± 0.000) | 0.999 (± 0.000) | $p < .001$ |
| Schaefer200 | 0.999 (± 0.001) | 0.999 (± 0.001) | 0.998 (± 0.002) | $p < .001$ |
| LAIRD | 0.809 (± 0.016) | 0.789 (± 0.019) | 0.857 (± 0.022) | $p < .001$ |
| Sliding-window size: 120s |  |  |  |  |
| AAL2 | 0.998 (± 0.001) | 0.999 (± 0.000) | 0.999 (± 0.001) | $p = .187$ |
| Schaefer200 | 0.999 (± 0.001) | 0.999 (± 0.001) | 0.999 (± 0.001) | $p = .195$ |
| LAIRD | 0.775 (± 0.018) | 0.814 (± 0.029) | 0.867 (± 0.017) | $p < .001$ |

*Note:* Classification performances were evaluated by using the area under curve (AUC). Higher AUC means better SVM classification performance. Each cell represents mean AUC ± SD. 'Permutation' means the permutation test results between the static-FC and the combined.

**Supplementary Table 4.**
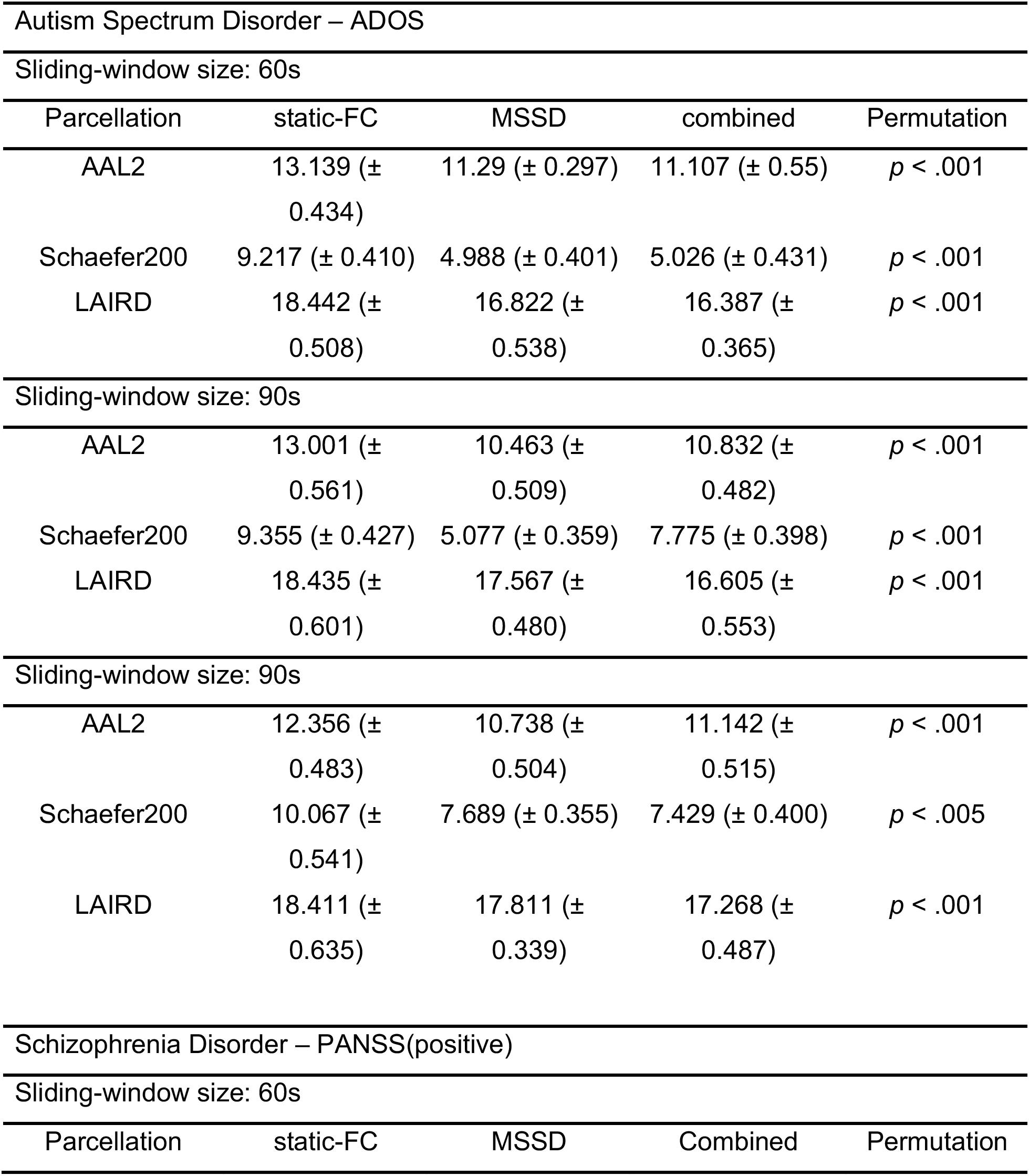
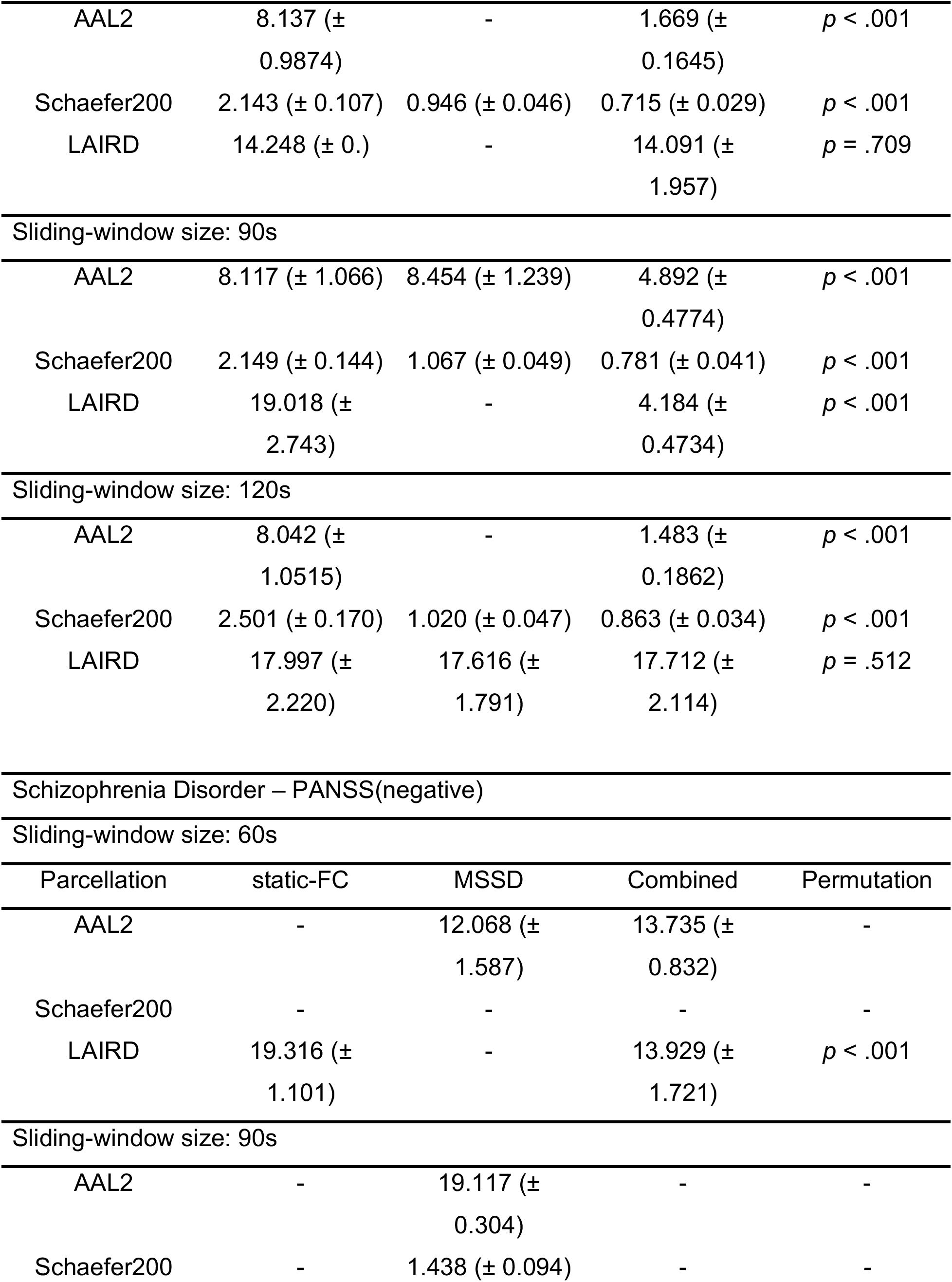
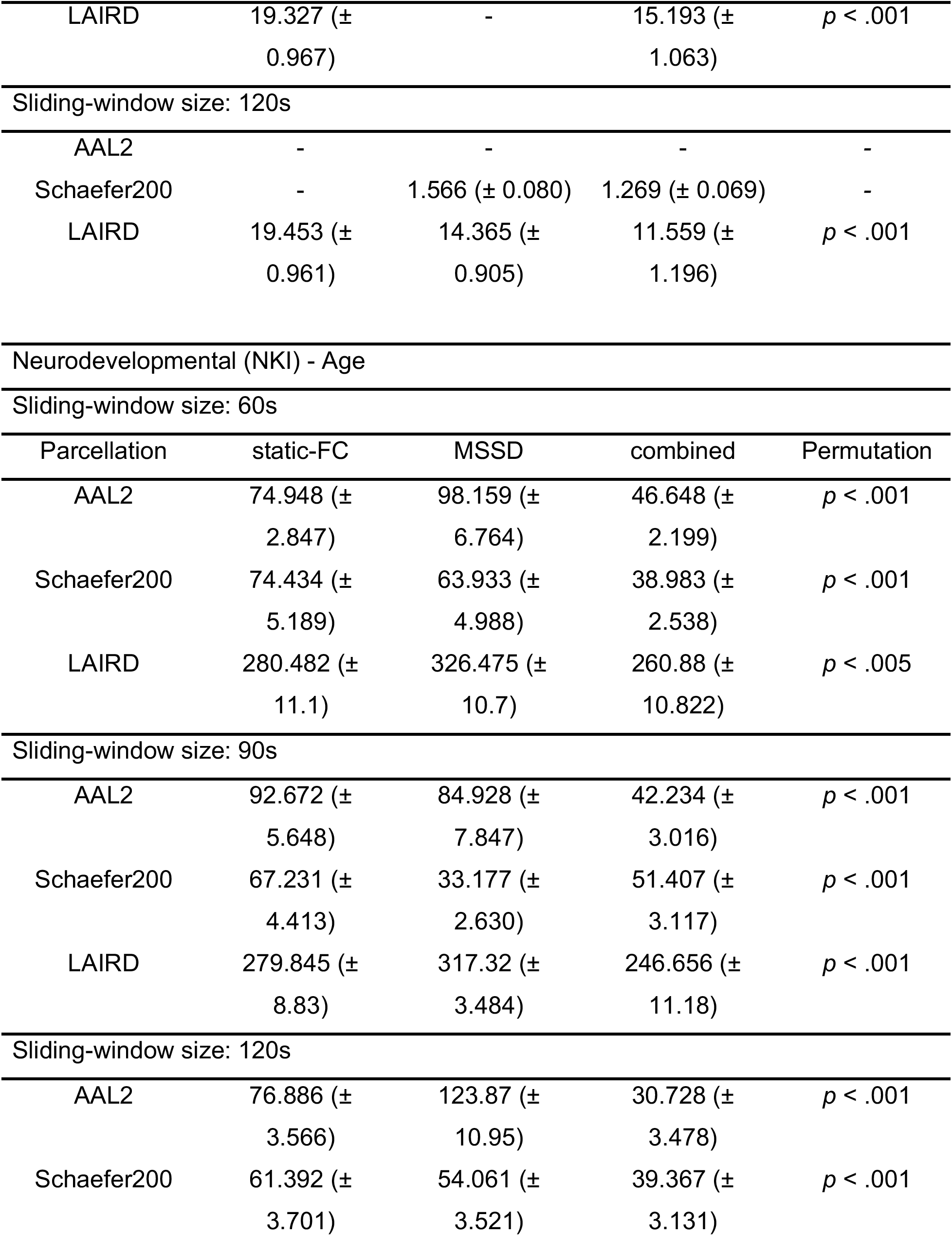
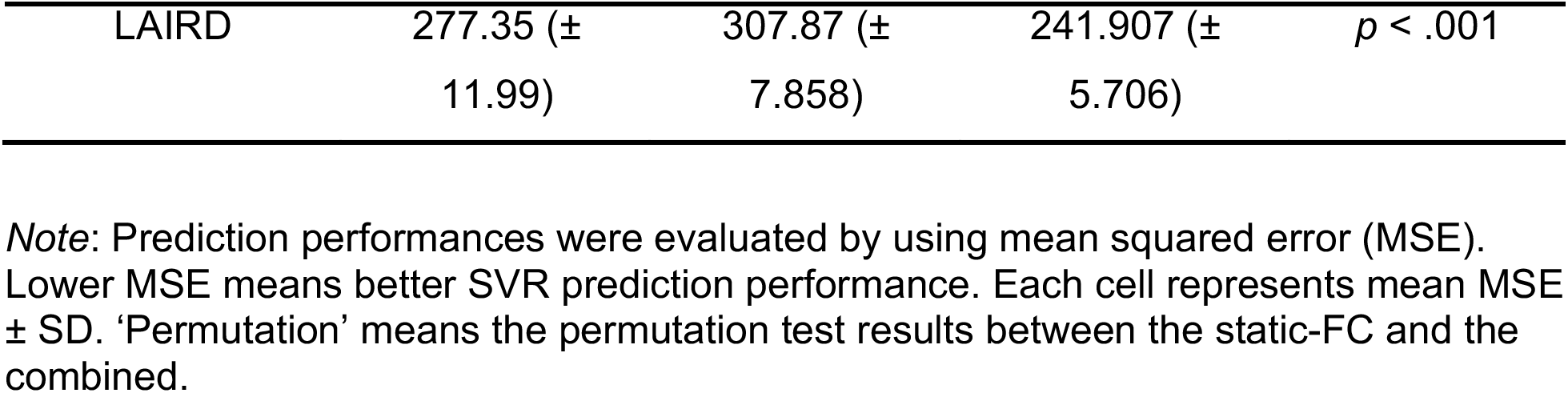
Predicting individuals’ characteristics performance with various sliding-window sizes (optimal lambda)

| Autism Spectrum Disorder – ADOS |  |  |  |  |
| --- | --- | --- | --- | --- |
| Sliding-window size: 60s |  |  |  |  |
| Parcellation | static-FC | MSSD | combined | Permutation |
| AAL2 | 13.139 (± 0.434) | 11.29 (± 0.297) | 11.107 (± 0.55) | $p < .001$ |
| Schaefer200 | 9.217 (± 0.410) | 4.988 (± 0.401) | 5.026 (± 0.431) | $p < .001$ |
| LAIRD | 18.442 (± 0.508) | 16.822 (± 0.538) | 16.387 (± 0.365) | $p < .001$ |
| Sliding-window size: 90s |  |  |  |  |
| AAL2 | 13.001 (± 0.561) | 10.463 (± 0.509) | 10.832 (± 0.482) | $p < .001$ |
| Schaefer200 | 9.355 (± 0.427) | 5.077 (± 0.359) | 7.775 (± 0.398) | $p < .001$ |
| LAIRD | 18.435 (± 0.601) | 17.567 (± 0.480) | 16.605 (± 0.553) | $p < .001$ |
| Sliding-window size: 90s |  |  |  |  |
| AAL2 | 12.356 (± 0.483) | 10.738 (± 0.504) | 11.142 (± 0.515) | $p < .001$ |
| Schaefer200 | 10.067 (± 0.541) | 7.689 (± 0.355) | 7.429 (± 0.400) | $p < .005$ |
| LAIRD | 18.411 (± 0.635) | 17.811 (± 0.339) | 17.268 (± 0.487) | $p < .001$ |
| Schizophrenia Disorder – PANSS(positive) |  |  |  |  |
| Sliding-window size: 60s |  |  |  |  |
| Parcellation | static-FC | MSSD | Combined | Permutation |
| AAL2 | 8.137 (± 0.9874) | - | 1.669 (± 0.1645) | $p < .001$ |
| Schaefer200 | 2.143 (± 0.107) | 0.946 (± 0.046) | 0.715 (± 0.029) | $p < .001$ |
| LAIRD | 14.248 (± 0.) | - | 14.091 (± 1.957) | $p = .709$ |
| Sliding-window size: 90s |  |  |  |  |
| AAL2 | 8.117 (± 1.066) | 8.454 (± 1.239) | 4.892 (± 0.4774) | $p < .001$ |
| Schaefer200 | 2.149 (± 0.144) | 1.067 (± 0.049) | 0.781 (± 0.041) | $p < .001$ |
| LAIRD | 19.018 (± 2.743) | - | 4.184 (± 0.4734) | $p < .001$ |
| Sliding-window size: 120s |  |  |  |  |
| AAL2 | 8.042 (± 1.0515) | - | 1.483 (± 0.1862) | $p < .001$ |
| Schaefer200 | 2.501 (± 0.170) | 1.020 (± 0.047) | 0.863 (± 0.034) | $p < .001$ |
| LAIRD | 17.997 (± 2.220) | 17.616 (± 1.791) | 17.712 (± 2.114) | $p = .512$ |

Schizophrenia Disorder – PANSS(negative) Sliding-window size: 60s
| Parcellation | static-FC | MSSD | Combined | Permutation |
| --- | --- | --- | --- | --- |
| AAL2 | - | 12.068 (± 1.587) | 13.735 (± 0.832) | - |
| Schaefer200 | - | - | - | - |
| LAIRD | 19.316 (± 1.101) | - | 13.929 (± 1.721) | $p < .001$ |
| Sliding-window size: 90s |  |  |  |  |
| AAL2 | - | 19.117 (± 0.304) | - | - |
| Schaefer200 | - | 1.438 (± 0.094) | - | - |
| LAIRD | 19.327 (±<br>0.967) | - | 15.193 (±<br>1.063) | $p < .001$ |
| Sliding-window size: 120s |  |  |  |  |
| AAL2 | - | - | - | - |
| Schaefer200 | - | 1.566 (± 0.080) | 1.269 (± 0.069) | - |
| LAIRD | 19.453 (±<br>0.961) | 14.365 (±<br>0.905) | 11.559 (±<br>1.196) | $p < .001$ |
| Neurodevelopmental (NKI) - Age |  |  |  |  |
| Sliding-window size: 60s |  |  |  |  |
| Parcellation | static-FC | MSSD | combined | Permutation |
| AAL2 | 74.948 (±<br>2.847) | 98.159 (±<br>6.764) | 46.648 (±<br>2.199) | $p < .001$ |
| Schaefer200 | 74.434 (±<br>5.189) | 63.933 (±<br>4.988) | 38.983 (±<br>2.538) | $p < .001$ |
| LAIRD | 280.482 (±<br>11.1) | 326.475 (±<br>10.7) | 260.88 (±<br>10.822) | $p < .005$ |
| Sliding-window size: 90s |  |  |  |  |
| AAL2 | 92.672 (±<br>5.648) | 84.928 (±<br>7.847) | 42.234 (±<br>3.016) | $p < .001$ |
| Schaefer200 | 67.231 (±<br>4.413) | 33.177 (±<br>2.630) | 51.407 (±<br>3.117) | $p < .001$ |
| LAIRD | 279.845 (±<br>8.83) | 317.32 (±<br>3.484) | 246.656 (±<br>11.18) | $p < .001$ |
| Sliding-window size: 120s |  |  |  |  |
| AAL2 | 76.886 (±<br>3.566) | 123.87 (±<br>10.95) | 30.728 (±<br>3.478) | $p < .001$ |
| Schaefer200 | 61.392 (±<br>3.701) | 54.061 (±<br>3.521) | 39.367 (±<br>3.131) | $p < .001$ |
| LAIRD | 277.35 (±<br>11.99) | 307.87 (±<br>7.858) | 241.907 (±<br>5.706) | $p < .001$ |

*Note:* Prediction performances were evaluated by using mean squared error (MSE). Lower MSE means better SVR prediction performance. Each cell represents mean MSE ± SD. 'Permutation' means the permutation test results between the static-FC and the combined.

## Footnotes

1 http://fcon_1000.projects.nitrc.org/indi/retro/cobre.html

2 http://fcon_1000.projects.nitrc.org/indi/abide

3 http://human.brain-map.org

